# TX-Phase: Secure Phasing of Private Genomes in a Trusted Execution Environment

**DOI:** 10.1101/2024.09.16.613301

**Authors:** Natnatee Dokmai, Kaiyuan Zhu, S. Cenk Sahinalp, Hyunghoon Cho

## Abstract

Genotype imputation servers enable researchers with limited resources to extract valuable insights from their data with enhanced accuracy and ease. However, the utility of these services is limited for those with sensitive study cohorts or those in restrictive regulatory environments due to data privacy concerns. Although privacy-preserving analysis tools have been developed to broaden access to these servers, none of the existing methods support haplotype phasing, a critical component of the imputation workflow. The complexity of phasing algorithms poses a significant challenge in maintaining practical performance under privacy constraints. Here, we introduce TX-Phase, a secure haplotype phasing method based on the framework of Trusted Execution Environments (TEEs). TX-Phase allows users’ private genomic data to be phased while ensuring data confidentiality and integrity of the computation. We introduce novel data-oblivious algorithmic techniques based on compressed reference panels and dynamic fixed-point arithmetic that comprehensively mitigate side-channel leakages in TEEs to provide robust protection of users’ genomic data throughout the analysis. Our experiments on a range of datasets from the UK Biobank and Haplotype Reference Consortium demonstrate the state-of-the-art phasing accuracy and practical runtimes of TX-Phase. Our work enables secure phasing of private genomes, opening access to large reference genomic datasets for a broader scientific community.

## Introduction

Genotype imputation servers [1] are invaluable tools for the genomics community, providing streamlined analysis pipelines that enhance the quality and resolution of genotype datasets. By utilizing reference panels of high-quality genomes, these servers perform *phasing* to estimate the haplotypes in the user’s data, followed by *imputation* to infer missing genotypes not captured in the original genotyping experiment. These servers are especially beneficial for users who lack access to extensive reference panels, sufficient computing resources, or expertise in the necessary data processing steps, enabling them to outsource these tasks to a service provider. However, despite their broad utility, the need to upload private genomic data to a remote server restricts access to these servers for users with particularly sensitive data (e.g., linked to rare health conditions or underrepresented populations) or those working in environments with stringent data sharing constraints stemming from privacy concerns.

The rapidly advancing field of privacy-enhancing technologies (PETs) offers a promising approach to expanding access to servers offering genome analysis as a service. This is achieved by enabling the secure processing of users’ data, without requiring complete trust in the service provider and their security precautions. For example, recent work [2] has shown that genotype imputation can be securely and accurately performed within Trusted Execution Environments (TEEs) [3]. TEEs are isolated computing environments equipped with hardware/microprocessor-level safeguards that maintain the confidentiality of user data and the integrity of computation, even from other entities with high-privileged access (e.g., administrative access) to the same device. While cryptographic tools based on homomorphic encryption have also been proposed to partially address the imputation workflow [4, 5], they face limitations in computational flexibility compared to TEEs, resulting in significant accuracy loss due to the algorithmic simplifications needed to achieve practical runtimes [2].

Unfortunately, none of the existing solutions based on PETs offer a viable secure alternative to existing imputation services, as they fail to support the critical task of phasing. Haplotype phasing [6–10] refers to the task of partitioning an individual’s diploid genotypes into a pair of haplotype sequences, one inherited from each parent. Since imputation algorithms rely on information about identity-by-descent (IBD) sequence sharing among individuals, which is characterized at the haplotype level, input data must be phased for accurate imputation. However, phasing, like imputation, is challenging for individual researchers to perform with limited data or computing resources, which has led to the widespread use of imputation servers for *both* phasing and imputation tasks. Moreover, phasing is becoming increasingly important as a standalone task, independently of imputation, for exploring various haplotype-level properties of the human genome [11, 12].

Developing a secure method for phasing poses significant challenges. The combinatorial nature of phasing, involving the assignment of alleles to haplotypes across many loci, inherently results in high computational costs, even without privacy considerations. Moreover, in the context of TEEs, the necessity to redesign algorithms to mitigate side-channel leakages [13], i.e., ensuring that memory access and timing patterns do not expose private information (referred to as *oblivious* computation), often introduces substantial computational overhead. This mitigation is essential because a side-channel leakage could be exploited to extract sensitive genomic data processed within a TEE [2, 14]. These limitations have, thus far, hindered the practical application of these technologies.

This work introduces a secure phasing method, TX-Phase, which overcomes these challenges to provide the first practical tool for reference-based phasing of private genomes in a remote computing environment. TX-Phase offers robust security based on TEEs, additionally incorporates a comprehensive mitigation of side-channel leakages, and obtains practical accuracy and runtime performance. This is achieved through our novel algorithmic techniques based on compressed reference panels and dynamic scaling of fixed-precision numbers (fixed points). Our results show that TX-Phase matches the accuracy of a state-of-the-art phasing algorithm (SHAPEIT4; [7]), even surpassing the accuracy of the tool currently used by imputation servers (Eagle2; [6]), all while ensuring data privacy. Combined with previous work on secure imputation [2], our work marks the completion of an end-to-end TEE-based solution that enhances the security of phasing and imputation services and broadens their use for a wider range of researchers. Furthermore, our techniques for oblivious computation on genome sequences could accelerate the development of other genomic analysis workflows to be securely provided as a service.

## Results

### Overview of TX-Phase

The main application setting of TX-Phase involves two entities: a user with unphased (diploid) genotype data and a service provider (SP) with a large haplotype reference panel. The user wishes to run a reference-based phasing algorithm on the user’s dataset with the SP’s reference panel. However, the reference panel in question is assumed to be too large for download or under strict access control, preventing the user from performing phasing locally and requiring the SP to perform the analysis on the user’s behalf. To support this service, SP could either use their own computing servers or provision third-party servers provided by commercial cloud services (e.g., Microsoft Azure). The key challenge addressed by our work is to enable such outsourcing of computation while keeping the user’s private data protected from other entities, including the SP, the operator of the computing server (could be the same as the SP), and adversaries who may obtain unauthorized access to the server. TX-Phase is designed to securely and efficiently support this workflow with state-of-the-art accuracy.

The security of TX-Phase is primarily based on the SP’s use of a Trusted Execution Environment (TEE) [3]. A TEE ensures that the user’s data is decrypted and analyzed only within an isolated computing environment on the remote server, called a *secure enclave*, which is protected from the rest of the system and other users (or operators) of the machine through secure hardware and cryptographic protocols. In addition to data confidentiality, most TEEs provide a mechanism for the user to verify the integrity of the secure enclave that was created to process the user’s workload, e.g., through a process called remote attestation [15]. While we demonstrate TX-Phase based on the Intel SGX framework [16], the most prominent TEE technology to date, our method can be easily deployed with other TEE technologies (e.g., Intel TDX [17] or AMD SEV [18]).

TX-Phase provides robust security beyond typical TEE applications by providing additional protection against side-channel leakages. A *side channel* [13] is an unintended pathway through which information about the user’s private data, held within a secure enclave, could be inferred. In particular, timing and memory side channels leak information through variations in operation runtimes or memory access patterns that depend on private data [19, 20]. This is an inherent limitation in SGX and other TEE architectures, which is often acknowledged but not explicitly addressed for the sake of computational performance. Previous work demonstrated that side channels in genomic analysis algorithms could be exploited to infer private genotypes even if the algorithm is running in a TEE [2, 14]. TX-Phase introduces algorithmic techniques to remove all known timing and memory side channels in the process, resulting in what we call an *oblivious phasing* algorithm, while maintaining practical performance.

We highlight two key algorithmic techniques introduced in TX-Phase for efficient oblivious phasing: *compressed haplotype selection* and *dynamic fixed-point arithmetic*. First, compressed haplotype selection leverages a compressed reference panel (based on the M3VCF representation [1]) to accelerate various components of the phasing algorithm by operating only on unique haplotype sequences within each genomic block. We apply this technique to substantially lower the costs of (1) querying the positional Burrows-Wheeler transform (PBWT) [21] data structure to find the reference haplotypes that are most related to the target sample and (2) filtering the reference panel to construct a conditioned panel used for phase estimation.

Next, dynamic fixed-point arithmetic eliminates the need for floating-point operations, which introduce timing side channels [22]. Fixed-point numbers, which represent fractional numbers using integers by multiplying them with a fixed scaling factor and then rounding, prevent side channels by relying only on constant-time integer operations, but suffer from limited precision. We address this limitation by introducing additional scaling factors shared among groups of numbers in matrices and vectors, which are dynamically adjusted throughout the analysis to maintain precision based on the magnitude of the data values. Our results show that this technique enables accurate probabilistic inference and sampling in our phasing algorithm. We provide further details of the TX-Phase algorithm and our techniques in Methods.

The following summarizes the workflow of TX-Phase. First, via a remote attestation protocol, the user verifies the TEE to ensure its security standing and TX-Phase’s program binary in a secure enclave to ensure it is not tampered with. Next, the user establishes a secure channel with the enclave to ensure sensitive inputs and outputs are not leaked or tampered with. The user then uploads input genotypes directly to the enclave via the secure channel. Subsequently, the SP securely phases the input genotypes with TX-Phase inside the enclave without exposing timing and memory side channels. Optionally, the SP performs imputation on the phased genotypes, also within the enclave, using a previously developed algorithm (e.g., SMac [2]). Finally, the user downloads the phased (and imputed) haplotypes from the enclave via the secure channel. We illustrate our workflow and key techniques in Figure 1.

**Fig. 1.**
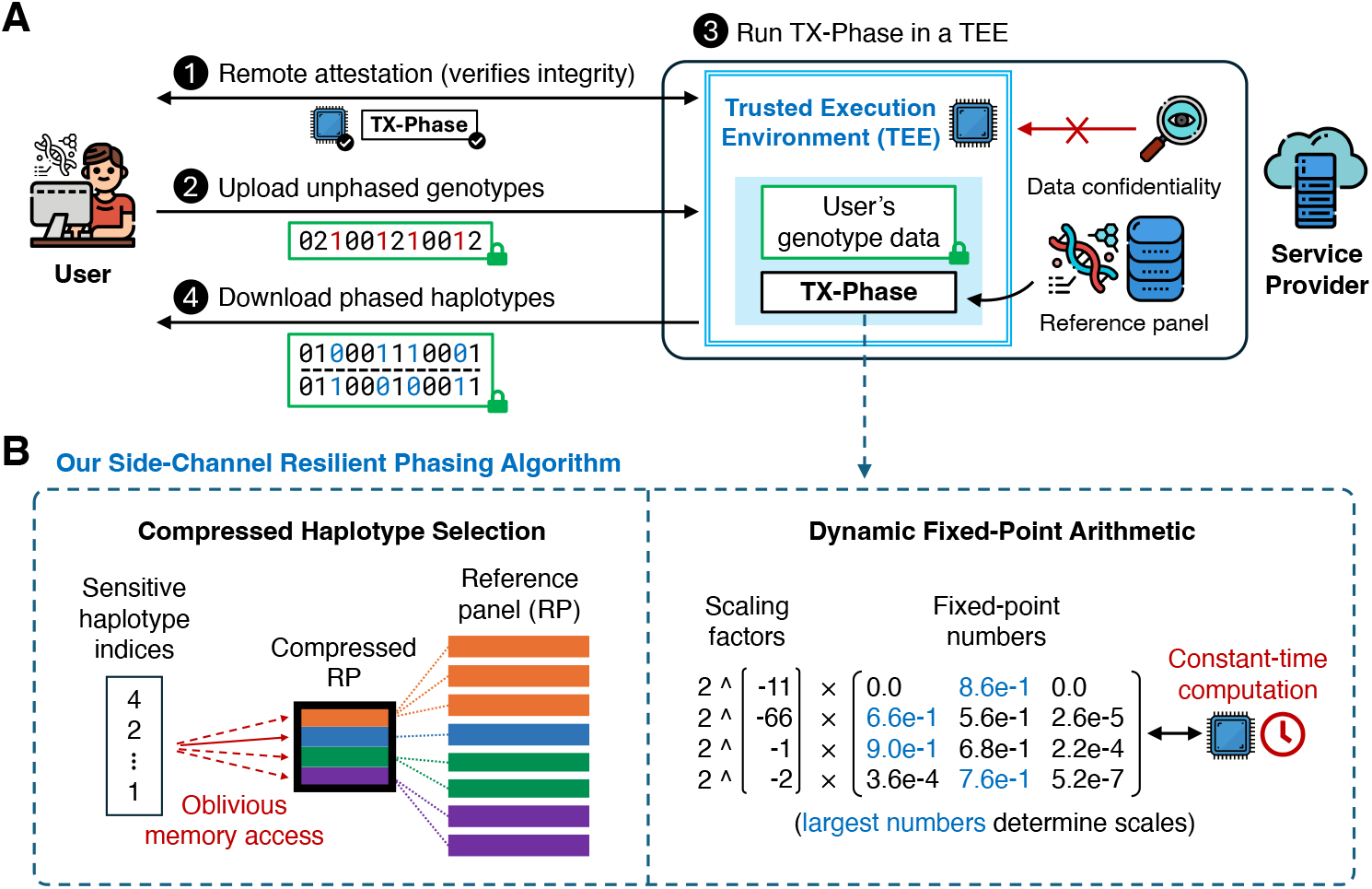
Overview of TX-Phase. We illustrate the workflow of TX-Phase (**A**) and our key algorithmic techniques (**B**). Our workflow proceeds as follows: (1) The user and the service provider’s Trusted Execution Environment (TEE; also called a secure enclave) engage in a remote attestation protocol to verify the integrity of the TEE and the TX-Phase program binary. (2) The user establishes a secure channel with the TEE and securely uploads unphased genotypes via the channel. (3) The service provider runs TX-Phase inside the secure enclave with a locally held haplotype reference panel to phase the user’s genotype data. (4) The user downloads the output phased haplotypes from the TEE via the secure channel. TX-Phase offers robust security, ensuring confidentiality of the user’s genomic data from the service provider and other users of the system. We introduce two key techniques—*compressed haplotype selection* and *dynamic fixed-point arithmetic*—which enabled us to design a phasing algorithm for TEEs that is efficient, accurate, and safeguarded against the risk of side-channel data leakage. Compressed haplotype selection refers to our novel use of compressed reference panels throughout the algorithm to minimize the overhead of memory-oblivious computation. Dynamic fixed-point arithmetic is our approach for enhancing the precision of fixed-point operations necessary for constant-time computations, achieved by dynamically adjusting the scales of numbers in matrices and vectors. Our results show that both techniques are essential for the secure and practical deployment of TEE-based haplotype phasing.

### TX-Phase provides state-of-the-art phasing accuracy

We first evaluate the phasing accuracy of TX-Phase on 424 parent-offspring trios from the UK Biobank (UKB) and compare with two state-of-the-art phasing tools: SHAPEIT4 [7] and Eagle2 [6]. TX-Phase is closely modeled after SHAPEIT4’s algorithm, which we comprehensively redesigned to achieve the side-channel resilience property for secure deployment in a TEE (Methods). Eagle2 represents the current method of choice in most existing imputation servers, including the Michigan and TOPMed imputation servers. We focused our analysis on Chromosome 20 (19,959 genetic variants) and sampled reference panels of varying sizes (20K, 40K, 100K, 200K, and 400K haplotypes) from the remaining UKB cohort. Phasing accuracy is measured in terms of *switch error rate* (SER), which refers to the proportion of consecutive heterozygous sites that are incorrectly phased compared to Mendelian phasing.

We observed nearly identical mean accuracies between TX-Phase and SHAPEIT4 across all reference panel sizes without any statistically significant difference (Figure 2A; two-sided Wilcoxon signed-rank test [WSR] *p >* 0.21). Due to the stochastic nature of the phasing algorithm, small differences in accuracy between the two methods are observed for individual samples (Figure 2B), but this difference was not skewed toward either method (e.g., WSR *p* = 0.50 for the 400K panel). Furthermore, TX-Phase significantly outperformed Eagle2 in accuracy (WSR *p <* 1.7 *×*10^*−*11^) for all but the smallest 20K reference panel (Figure 2A and C). This suggests that TX-Phase could improve both the security and the accuracy of the current phasing workflow on existing imputation servers.

**Fig. 2.**
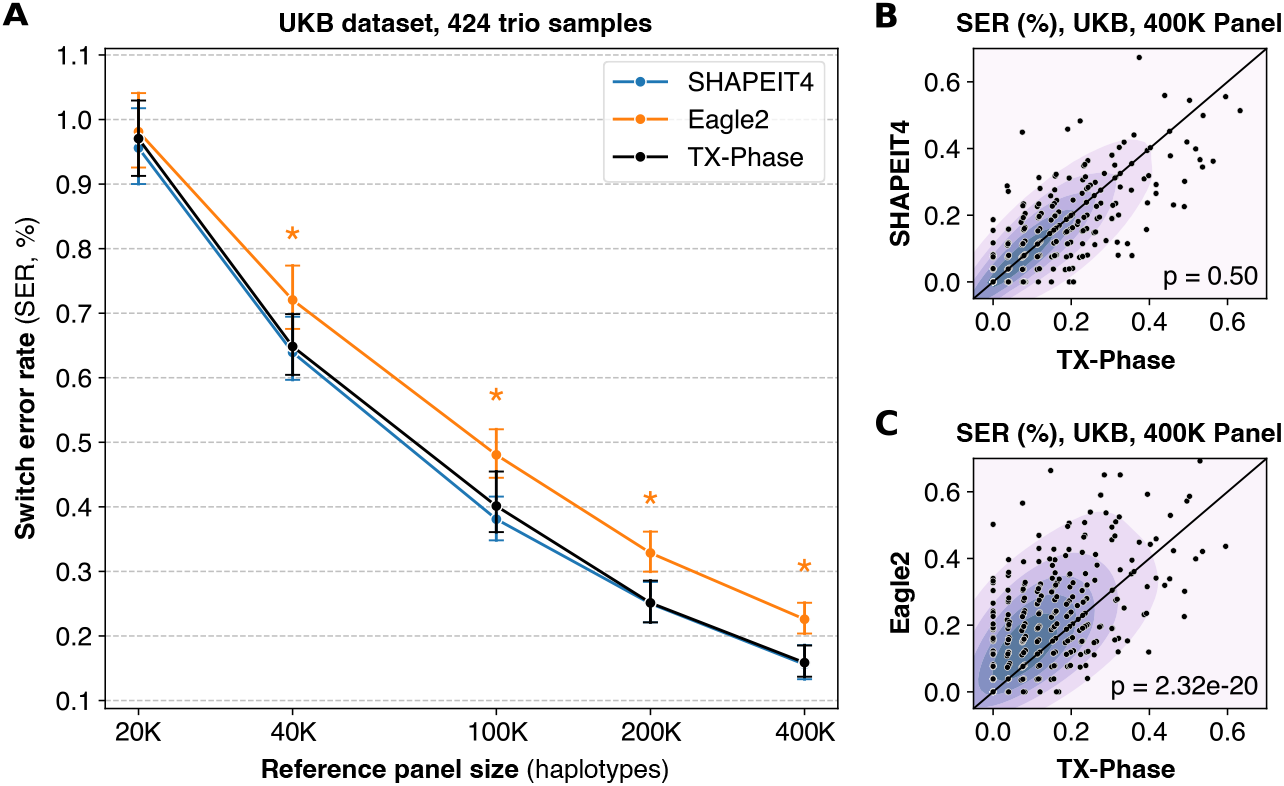
TX-Phase obtains state-of-the-art phasing accuracy on the UK Biobank dataset. **(A)**We compared the accuracy of TX-Phase with two standard phasing methods, SHAPEIT4 [7] and Eagle2 [6], on 424 trio samples from the UK Biobank (genotyping array data, Chromosome 20) using subsets of the remaining cohort of varying sizes as the reference panel. Accuracy is measured by switch error rates (SERs) with trio-based phasing as the ground truth. Markers indicate the mean SERs and error bars represent 95% confidence intervals. Asterisks denote highly significant differences (two-sided Wilcoxon signed-rank test, *p <* 1 *×*10^*−*10^)) between TX-Phase and Eagle2. The same statistical tests were performed between TX-Phase and SHAPEIT4, which did not result in any significant difference. We also compare the per-sample SERs of TX-Phase against those of SHAPEIT4 (**B**) and Eagle2 (**C**) for the 400K reference panel. The background heatmap visualizes marker density. For clarity, the display region is limited to [0, 0.7] for both axes. We show the p-values from the two-sided Wilcoxon signed-rank test, which suggest that TX-Phase significantly outperforms Eagle2 while obtaining comparable accuracy with SHAPEIT4. Consistent results are observed when phasing a sample from the Genome-in-a-Bottle Consortium using the 1000 Genomes and Haplotype Reference Consortium reference panels (Supplementary Figure 1).

For additional validation, we evaluated TX-Phase on the Ashkenazi trio from the Genome-in-a-Bottle (GIAB) Consortium dataset using the 1000 Genomes Phase 3 (1KG) dataset (5,008 haplotypes from 2,504 subjects) and the Haplotype Reference Consortium (HRC) dataset (54,330 haplotypes from 27,165 subjects) as reference panels (Methods). Consistent with the UKB results, TX-Phase achieved accuracy comparable to that of SHAPEIT4 and outperformed Eagle2 for both 1KG and HRC reference panels (Supplementary Figure 1). These results demonstrate that TX-Phase provides state-of-the-art phasing accuracy despite its reliance on fixed-point arithmetic due to the security constraints.

### TX-Phase is practical in running time and memory usage

We next evaluate the computational costs of TX-Phase with respect to both running time and peak memory usage and compare them with SHAPEIT4 and Eagle2. Due to the additional computational costs introduced by executing a program in a TEE and our side-channel protection techniques, some slowdown is expected for TX-Phase compared to the existing non-TEE methods. Nevertheless, we observed practical running times for TX-Phase on the UKB dataset across all reference panel sizes (Figure 3A). For example, TX-Phase took two minutes (123 secs) to phase Chromosome 20 using the 400K reference panel, reflecting an increase of 1.3x and 2.3x in running time compared to SHAPEIT4 (93 secs) and Eagle2 (53 secs), respectively. Extrapolating these results, we estimate that phasing a whole genome sample (e.g., with 800K variants) with the 400K haplotype reference panel would take 82 minutes using TX-Phase, compared to 63 minutes using SHAPEIT4 and 35 minutes using Eagle2.

**Fig. 3.**
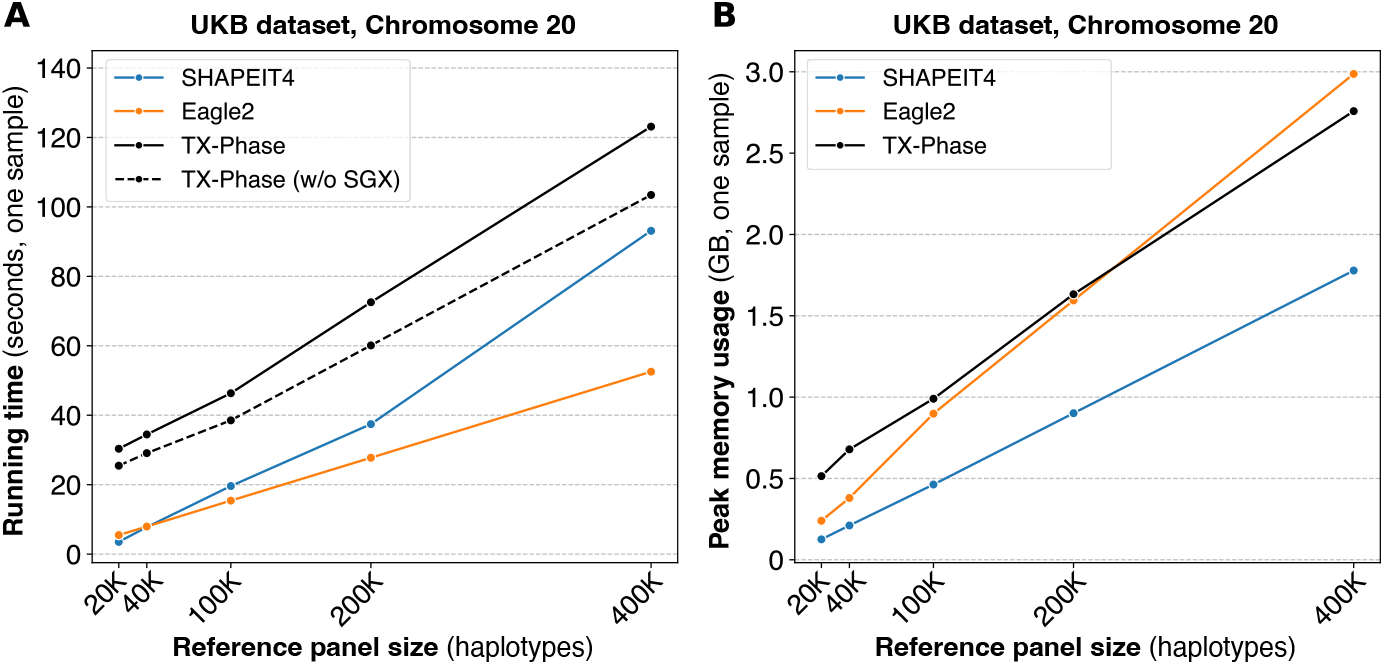
TX-Phase maintains practical running times for phasing individual samples while providing enhanced security. We report the running times (**A**) and peak memory usages (**B**) of TX-Phase, SHAPEIT4, and Eagle2 for phasing a sample in the UK Biobank (UKB) genotyping array dataset (Chromosome 20, 19,959 variants) with varying reference panel sizes up to 400K haplotypes. Additionally, we plot the running times of TX-Phase without SGX for comparison, where our program is executed in a conventional (unprotected) computing environment. The results demonstrate that TX-Phase effectively minimizes the overhead of side-channel resilient computation in TEEs, achieving practical performance comparable to existing tools, or only marginally more costly (e.g., by less than a factor of 3) on large reference panels.

To quantify the overhead of TEEs, we also timed the execution of TX-Phase outside the secure enclave for comparison. We observed small additional runtime costs for TX-Phase, e.g., 19% on the largest reference panel (Figure 3A). In comparison, running SHAPEIT4 or Eagle2 within the enclave through the Gramine framework [23] resulted in larger overhead costs of 71% and 43% on the largest panel, respectively (Supplementary Figure 2). Notably, TX-Phase was even faster than SHAPEIT4 on the largest panel when both were executed in a TEE, illustrating the effective design of TX-Phase that minimizes the TEE overhead. We further note that directly deploying existing phasing tools in a TEE introduces side-channel vulnerabilities and thus is not recommended; TX-Phase represents a practical alternative with robust protection against such risks.

The peak memory usage of TX-Phase was comparable to Eagle2 and 1.6 times that of SHAPEIT4 for the largest reference panel (Figure 3B). TX-Phase’s memory usage remained below 3 GB in all settings, which allows it to be deployed on low-cost servers with standard memory resources. Note that phasing the whole genome does not increase the memory usage of TX-Phase because fixed-size genomic windows are processed independently, which leads to memory usage that depends primarily on the size of the window and the number of haplotypes in the reference panel.

### Compressed haplotype selection enables scalable performance

TX-Phase introduces novel algorithmic techniques based on compressed reference panels to minimize the cost of oblivious computation in phasing. To demonstrate the significance of these techniques, we compared TX-Phase with its modified version that uses the original (uncompressed) reference panel following the same approach as the original SHAPEIT4 algorithm but with side-channel protection.

We focus on two analysis steps in the selection of conditional haplotypes (i.e., reference haplotypes that are closest to the phasing estimates of the target sample) that pose a computational bottleneck given a large reference panel: *PBWT querying* and *haplotype filtering*. PBWT querying refers to the task of retrieving the indices of reference haplotypes with the longest suffix matches with the target sequence at different sequence positions using the PBWT data structure. Haplotype filtering represents the subsequent step of constructing a reduced reference panel consisting only of the chosen haplotypes for use in the hidden Markov model (HMM)-based inference algorithm for phase estimation (Methods). TX-Phase accelerates both these steps using a compressed reference panel, which allows us to operate over the space of unique haplotypes within each genomic block and avoids full scans over the original panel, which is overwhelmingly costly when performed obliviously. For example, accessing a single haplotype in the panel could require redundantly accessing all haplotypes to hide the access index (Methods).

Our results show that our compression techniques are pivotal to TX-Phase’s practical running times (Figure 4). Based on the largest panel of 400K haplotypes, TX-Phase is more than 35 times faster than the alternative approach without our compression techniques for PBWT querying and 7 times faster for haplotype filtering, which together lead to a 15 times faster running time for the entire phasing pipeline. For whole-genome phasing (with 800K variants) using the 400K reference panel, we estimate that TX-Phase would result in a running time of 1.4 hours compared to 21 hours without our compression techniques.

**Fig. 4.**
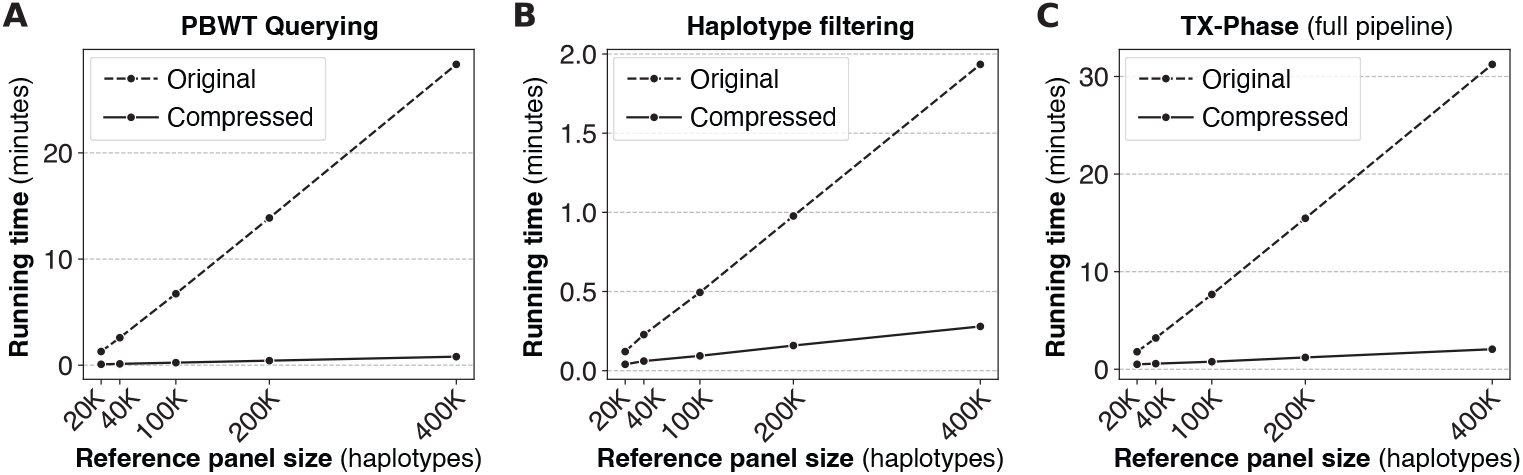
Our compressed haplotype selection techniques are essential for TX-Phase’s computational efficiency. We compare TX-Phase (Compressed) with an alternative side-channel resilient implementation without compression (Original), which more closely follows SHAPEIT4’s approach, with respect to their running times. We measured the running time for phasing one sample (Chromosome 20) in the UK Biobank dataset across varying reference panel sizes. The results are summarized for different components of the algorithm, including PBWT querying (**A**) and haplotype filtering (**B**), as well as the full phasing pipeline (**C**). Our compressed algorithms substantially reduce the running time of TX-Phase, which otherwise becomes prohibitive for large reference panels.

### Dynamic fixed-point arithmetic ensures accurate phasing

We designed TX-Phase to use only constant-time, fixed-point arithmetic operations to circumvent the timing side channels introduced by floating-point operations. However, since fixed-point numbers represent a fixed range of values with limited precision, without careful algorithm design, they could introduce a substantial accuracy loss in probabilistic calculations that are sensitive to small values.

Previous work on TEE-based imputation [2] proposed performing probability calculations in the logarithmic domain to minimize precision loss at the cost of extra computation required for addition and subtraction of probabilities, which become non-linear operations in the log space. Although this approach was shown to be effective for the imputation task, we observed that the same approach would lead to prohibitive running times for phasing due to the larger dimensions of the matrices involved, which introduce more additions relative to other operations. Our complexity analysis suggests that adopting the log-domain approach would result in a 100x slowdown of TX-Phase performance (Supplementary Table 1). To address this limitation, TX-Phase incorporates the novel approach of representing the original probabilities as fixed-point numbers, without log transformation, but with dynamically updated scaling factors for groups of numbers (e.g., rows or columns of a matrix; see Methods).

Comparing TX-Phase with an alternative implementation without our dynamic scaling technique revealed that our approach substantially reduces the SER on the UKB dataset, consistently across different reference panel sizes (Figure 5). For instance, with the 400K reference panel, TX-Phase achieves an average SER of 0.16%, compared to 0.55% without our technique, representing an improvement of more than threefold. Furthermore, our technique allows TX-Phase to closely match the accuracy of SHAPEIT4, despite TX-Phase’s reliance on fixed-point numbers with limited precision. The accuracy loss without dynamic scaling typically results from integer underflows, where entire distributions are incorrectly flattened to a uniform distribution due to small probabilities turning into zeros, thus introducing problems in the sampling process of the phasing algorithm.

**Fig. 5.**
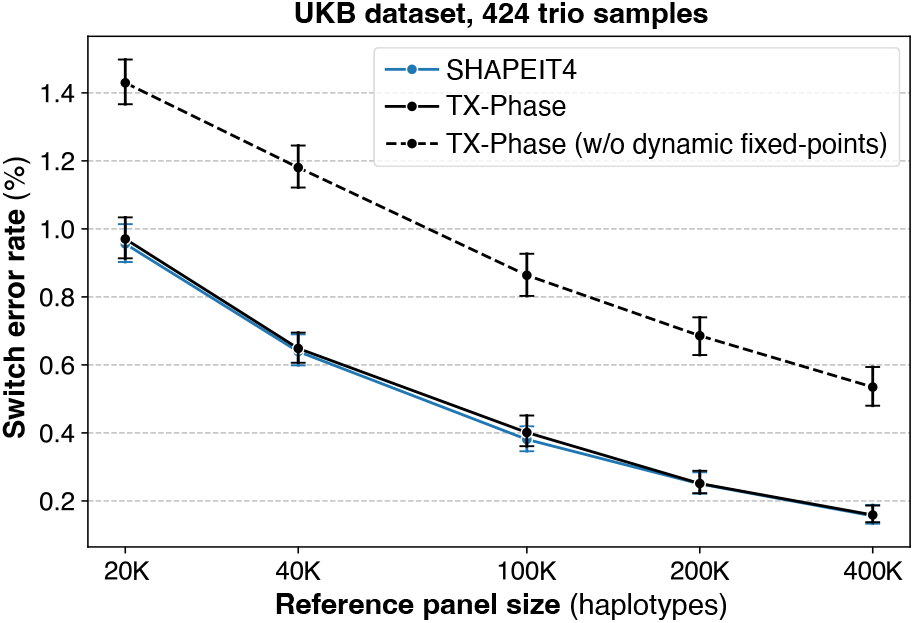
Our dynamic fixed-point arithmetic techniques ensure accurate phasing of TX-Phase under precision constraints. We compare the phasing accuracy of TX-Phase, measured by the switch error rate (SER), with an alternative implementation that lacks our dynamic scaling techniques for fixed-point matrices and vectors. The evaluation is conducted on 424 trio samples (Chromosome 20) from the UK Biobank (UKB) dataset, using reference panels of varying sizes from the remaining cohort. Markers indicate the mean SERs and error bars represent 95% confidence intervals. We also include the accuracies of SHAPEIT4 for comparison, which is shown to be comparable with TX-Phase. Without our dynamic fixed-point techniques, the accuracy of TX-Phase significantly deteriorates.

## Discussion

Our method TX-Phase demonstrates the practicality of secure haplotype phasing powered by TEE technologies, with state-of-the-art phasing accuracy and robust side-channel protection. Together with an existing tool for imputation [2], our work represents the completion of an end-to-end TEE-based solution for genotype imputation servers, encompassing both phasing and imputation workflows. Our approach complements and strengthens existing security measures by minimizing the trust required in the security of the remote computing infrastructure. The enhanced security could broaden access to these services for users in restrictive environments, for example by facilitating compliance with regulations such as HIPAA [24], GDPR [25], and Germany’s Patient Data Protection Act (PDSG) [26].

The financial cost of deploying a TEE application in the cloud environment has drastically decreased in recent years to the same level as traditional computing. For example, a Microsoft Azure instance in the DCsv3 series, equipped with Intel SGX version 2, is only about 50% more expensive for the hourly rate compared to the standard Ev5 series with equivalent specifications, according to the Azure price calculator (https://azure.microsoft.com/en-us/pricing/calculator). Therefore, deploying TX-Phase at scale with a large number of CPU cores and multiple virtual machine instances is economically viable. Our current estimate of the cost of phasing a whole genome sample with 800K variants using a reference panel of 400K haplotypes with TX-Phase is $0.26 (on Microsoft Azure DC2sv3), compared to phasing with SHAPEIT4 at $0.13 and Eagle2 at $0.07 (both on Microsoft Azure E2v5).

Although our experiments focused on single-sample phasing for comparison, processing an input dataset with multiple samples can be easily supported via parallelization. We confirmed that the overhead associated with such parallelization is minimal for TX-Phase (Supplementary Figure 3). SHAPEIT4 incorporates additional optimizations to process multiple samples more efficiently; similarly extending our approach to minimize redundant computation across samples, e.g., by sharing the compressed PBWT data structure, is a meaningful direction for future work. Furthermore, our method can be extended to include rare variants in the output haplotypes following the approach introduced in SHAPEIT5 [8].

Another future direction is to explore general applications of the algorithmic techniques introduced in TX-Phase. Our compressed PBWT data structure and associated operations could accelerate PBWT-based sequence search in large genomic databases in analysis tasks beyond the TEE setting, including other phasing methods [6, 9], imputation [10], and identity-by-descent (IBD) detection [27]. Moreover, our techniques for efficient and accurate oblivious computation could accelerate the development of other TEE-based tools to enable confidential analysis of sensitive biomedical datasets.

## Methods

### Review of Intel SGX Trusted Execution Environment (TEE)

Intel Software Guard Extensions (SGX) [16] is one of the most mature TEE platforms for user-level applications, having been in production since 2015 when the first SGX-equipped CPU was introduced to the market. SGX version 2 [28], the most recent iteration of the SGX architecture, is integrated into modern server-grade Intel Xeon CPUs to support confidential computing in remote environments. SGX version 2 introduces the Enclave Dynamic Memory Management (EDMM) technology, which enables a substantially larger protected memory region and higher memory throughput for large-scale applications.

Intel SGX employs hardware root-of-trust, cryptographic techniques, and decentralized trust to ensure the confidentiality and integrity of data and code within a secure enclave. Confidentiality ensures that no unauthorized processes can access protected information inside the enclave during runtime. Integrity guarantees that architectural enclaves (i.e., system enclaves facilitating SGX functionality), Trusted Computing Base (TCB) and libraries signed by Intel, and data and codebases verified by the remote user cannot be modified by unauthorized processes during runtime. This allows sensitive applications to be deployed in untrusted computing environments, even where the operating system might be compromised. Remote users can verify the security status of SGX hardware and the authenticity of runtime applications via a process called remote attestation [15], and deny participation before uploading sensitive input if the security status is not up to standard, for example, if there are unmitigated vulnerabilities.

Despite its promise, applying this technology in practice requires careful consideration of the security it provides and the implementation of complementary security safeguards. The research community has discovered a variety of security vulnerabilities in its architecture over the years [19, 20]. Although most security vulnerabilities have been patched in the latest generation of Intel CPUs, side-channel leakage issues are left to method developers to address through software mitigations. These vulnerabilities highlight the trade-off between security and efficiency in modern CPU architecture, making them unlikely to be entirely eliminated in the design of TEEs beyond SGX. For instance, many side channels between isolated processes stem from shared CPU resources such as CPU caches and computing units such as the arithmetic logic unit (ALU), which can be exploited to extract sensitive information.

Our key approach in TX-Phase is to eliminate the possibility of these types of leakage at the algorithmic level through data-oblivious computation, where control flow, timing, and memory access patterns are entirely independent of sensitive data.

### Overview of TX-Phase Workflow and Security Model

TX-Phase ‘s workflow involves two primary participants: the service provider (SP) and the user. The SP deploys TX-Phase within a secure enclave in a TEE-enabled computing environment. The user verifies the enclave and submits unphased genotypes as input samples to the SP. The SP then uses TX-Phase, alongside a large reference panel of haplotype sequences (which is accessible only to the SP), to phase the genotypes and returns the phased genotypes to the user.

We consider the setting where the SP has complete control over the computing environment, including the highest privileged access to the operating system and physical devices. This encompasses both common scenarios where the SP manages their own computing infrastructure and where the service is hosted on a virtual machine in a commercial cloud platform.

Based on TEEs, TX-Phase provides robust protection of the user’s input data against a potential attacker with the same level of access as the SP. The attacker aims to extract the user’s genotypes by modifying or inspecting various aspects of the system while the server processes the user’s workload. In some TEE platforms with integrity protection, such as Intel SGX and Intel TDX, the attacker may also attempt to compromise the integrity of the phasing results.

Importantly, we assume that the SP is acting in good faith and is motivated to prevent such an attacker from succeeding. To this end, the SP may have implemented other security measures, such as access control, scheduled data deletion, and regular software updates. Our TEE-based method should be viewed as a complementary tool to significantly enhance the security offered by these services. In essence, TEE allows the SP and the user to collaborate to achieve the best possible security outcomes.

In the following, we assume that the TEE-enabled computing environment’s security status is current and focus on our mitigation of side-channel vulnerabilities. For Intel SGX, the user can verify the security configuration of the SGX server via remote attestation and may choose not to participate if the configuration is not satisfactory. This ensures that the sensitive input is not transmitted to the server before verification.

### Side-Channel Vulnerabilities in Haplotype Phasing

The following are two main categories of side-channel vulnerabilities in TEEs demonstrated in the literature. As described in later sections, TX-Phase provides comprehensive protection against both types of vulnerabilities.

#### Timing side channels

Timing information resulting from the computation of sensitive data can be exploited through side channels to infer the sensitive data. In phasing tools, one prominent example is the *emission code gadget* in the hidden Markov model (HMM) used to model each haplotype as a mosaic of reference panel haplotypes [7]. The emission probability value, used to update beliefs over the hidden states in the HMM, is determined by whether the user’s genotype matches the reference genotype at a given position. If one value is large and the other is small (subnormal) in the floating-point representation, the floating-point multiplications between these two branches can take different amounts of time [22]. This discrepancy can be exploited by side-channel attacks, such as port contention [2, 29], to infer which branch has been executed and thus the sensitive genotype.

#### Memory-access side channels

The computation of sensitive data can lead to unique memory access patterns, which an external process can observe via side channels to infer the sensitive data. In phasing tools, one such pattern occurs when haplotypes in the reference panel are searched based on their similarity to the user’s input genotypes to construct a conditioned reference panel. Knowing that certain haplotypes have been looked up may reveal their similarity to the user’s input genotypes. Techniques such as Prime+Probe [14] have been demonstrated to exploit this class of vulnerabilities.

### Our Side-Channel Mitigation Strategies

Many software countermeasures to side channels in TEE have been proposed in the literature [20], each with various trade-offs between security, performance, and ease of use. TX-Phase not only builds upon these general strategies, but also introduces new algorithmic techniques to obtain a scalable solution for TEE-based phasing with comprehensive side-channel mitigation. We first outline these techniques and describe them further after presenting the phasing algorithm in more detail.

#### Memory-oblivious algorithms

TX-Phase’s phasing method requires memory access that is dependent upon sensitive genotypes, particularly during the construction of conditioned reference panels, where the user’s sensitive input genotypes are used to select and access haplotypes in the reference panel. Thus, an attacker who observes the memory access patterns could infer the user’s genotypes, which necessitates the use of oblivious computation approaches whereby the operations are modified to keep the memory access the same regardless of the sensitive data.

For example, to look up elements in an array without leaking the secret index, we employ oblivious RAM (ORAM) algorithms [30]. ORAM introduces redundant memory access to obfuscate true access patterns. ORAM algorithms typically have an asymptotic runtime of *O*(log *N*) to look up an item in an array of size *N*. However, in practice, there is substantial concrete overhead in runtime and storage, making many ORAM techniques suitable primarily for accessing larger arrays and databases than those encountered in phasing. Additionally, standard ORAMs do not distinguish read and write operations, leading to the two having the same cost, which is not suitable for read-heavy applications like ours.

For these reasons, TX-Phase uses a simple *linear scanning* ORAM for array access, which hides memory access patterns by pretending to access all items in an array. Despite its suboptimal asymptotic complexity of *O*(*N*), its simplicity and lack of runtime/storage overhead from internal data structures make it the most efficient option for moderately sized arrays. In addition, the resulting separation of read and write operations in TX-Phase enables the use of fast sequential reads in modern CPU architectures. Moreover, we make a standard assumption that no attacks can distinguish memory access within a 64-byte cache line (which comes with a few requirements, such as deactivating hyperthreading in Intel SGX [19, 31]). This allows our linear scanning ORAM to access every cache line instead of every array item, thus further improving its speed. For sorting and filtering operations, TX-Phase uses a combination of oblivious techniques based on Bitonic sort [32], bubble sort, and linear scanning ORAM.

Despite these efficient operations, memory-oblivious computation typically leads to an overwhelming computational overhead when large datasets are involved (e.g., the reference panel in phasing). TX-Phase introduces compressed data structures and associated operations for haplotypes, which are crucial to minimizing overhead and maintaining practical runtimes (see Accelerating Oblivious Phasing via Compressed Haplotype Selection).

#### Deterministic control flow

To ensure that the control flow of TX-Phase (such as if or for statements) is not affected by the sensitive input, we make the control flow fully deterministic and input-independent. For example, we ensure that an if statement always executes all possible branches, and for loops always continue for a fixed number of iterations. To minimize redundancy in conditional branches, this approach often requires that the algorithm be restructured to better separate computation on sensitive data from that on non-sensitive data.

#### Constant-time instructions

Certain CPU instructions in phasing algorithms are known to be input-dependent in running time, particularly floating-point arithmetic and integer division [22]. To mitigate this timing discrepancy, we follow Andrysco et al. [22] in using only constant-time CPU instructions for TX-Phase’s implementation of code involving sensitive data. This includes using fixed-point primitives (based on integers, whose running time is input-independent) to represent real numbers and implementing constant-time integer long division using bitwise operations.

In contrast to a previous work on genotype imputation [2], which proposed performing fixed-point arithmetic in log space for precision, TX-Phase develops a new framework based on dynamically scaled fixed-point numbers, described in Dynamic Fixed-Point Arithmetic for Accurate Constant-Time Computation, which allows TX-Phase to obtain state-of-the-art phasing accuracy.

#### Secure typing of sensitive data

Following the previous work on imputation [2], we adopt a secure typing system in Rust programming language to ensure that all sensitive input data is appropriately protected in TX-Phase. This is achieved by creating a wrapper around standard types and overriding their input-dependent subroutines. The secure types enforce a set of rules where: (1) any subroutines and computation involving them must be carefully implemented to be constant-time and data-oblivious in memory access, (2) computation of secure types must always result in a secure type, and (3) exposure of secure types into non-secure types must be explicit. The benefit of this approach is that any violation of the security rules can be detected at the time of compilation, thus speeding up software development and making the resulting code more reliable. In our work, we extend the secure types from the previous work to include secret indices that can only be used to access arrays via ORAM or other data-oblivious algorithms, which are used extensively throughout our phasing algorithm.

### TX-Phase’s Reference-Based Phasing Algorithm

TX-Phase adopts the reference-based phasing approach of a state-of-the-art tool SHAPEIT4 [7], which we thoroughly redesigned to remove all known timing and memory access side channels while maintaining efficient and accurate performance. Here, we provide an overview of the phasing algorithm before highlighting some key transformations introduced in TX-Phase.

TX-Phase utilizes the Markov Chain Monte Carlo (MCMC) algorithm to progressively improve the estimated phased haplotypes of the target sample through iterative sampling. In each iteration, a small but targeted *conditioned reference panel* is constructed, which is a subset of haplotypes from the original reference panel that share the longest identity-by-state (IBS) segments with the current estimate of the phased haplotypes. This haplotype subset is chosen efficiently using the positional Burrows-Wheeler transform (PBWT) data structure [21], enabling efficient searches for sequences with the longest suffix match with the target sequence at different sequence positions.

Given the conditioned panel, TX-Phase uses a variant of the Li-Stephens Hidden Markov Model (HMM) [33], which models each haplotype sequence as a mosaic of the reference haplotypes, to compute the transition probabilities over the phased genotypes between adjacent genomic segments, representing the current belief over the target phases. These probabilities are then used to sample a new pair of phased haplotype estimates, provided as the input to the next iteration. Following SHAPEIT4, the state space of the HMM in TX-Phase is augmented to include both the reference panel and the segmented haplotype model of the target sample. The latter represents all possible phased haplotypes as paths through a *genotype graph*, which includes short non-overlapping haplotype segments as nodes. Each segment covers a small number of heterozygous sites (three by default) to allow the enumeration of all possible phases of each segment for inference.

The MCMC algorithm for phasing proceeds through several different stages of the sampling process: (i) *Initialization*, where a heuristic strategy is used to guess the initial phases of the target sample to use as an initial search point; (ii) *Burn-in*, where some number of sampling iterations are performed to move the initial estimate closer to a desired sample from the posterior distribution over the phases; (iii) *Pruning*, where adjacent phased haplotype segments with high confidence are merged to reduce the search space; and (iv) *Main*, where the final, maximum-likelihood phases are determined via the Viterbi posterior decoding algorithm. TX-Phase inherits the default sampling procedure from SHAPEIT4, involving multiple rounds of alternating burn-in and pruning stages before the main iterations are performed to obtain the final phasing output.

### Accelerating Oblivious Phasing via Compressed Haplotype Selection

The size of the reference panels is a primary performance bottleneck for the phasing tools, affecting both the memory usage and the running time of algorithms. Typical panel sizes can range from tens of thousands of haplotypes (e.g., the Haplotype Reference Consortium dataset) to hundreds of thousands in the case of large genomic biobanks (e.g., the UK Biobank). Leveraging large panels is particularly challenging for TEE-based phasing because side-channel resilient algorithms result in conservative, worst-case runtime and introduce high redundancy for a task that would be instantaneous otherwise. For example, the cost of privately accessing a single haplotype in the reference panel using an ORAM approach (see Our Side-Channel Mitigation Strategies) can be *O*(log *N*) or *O*(*N*), where *N* is the number of haplotypes, compared to *O*(1) in the non-oblivious setting.

To improve the efficiency of oblivious phasing, we draw inspiration from fast imputation methods [1] to use a *compressed* reference panel, which can reduce its operational size by two orders of magnitude for large reference panels (e.g., the Haplotype Reference Consortium and UK Biobank dataset). Our compression method, based on the M3VCF encoding [1], splits the original reference panel into short contiguous genomic “blocks”. The haplotypes within each block are compressed to keep only the *unique* haplotypes observed. To preserve the ability to reconstruct the original sequences from each block, index maps are maintained to link the original samples with their corresponding unique haplotypes.

As described in the previous section, our phasing algorithm applies PBWT [21] to the reference panel to perform an efficient search for the “nearest neighbors” of the estimated target haplotypes based on the longest common suffix (LCS). Note that the neighbor search is performed at randomly sampled genomic positions, which introduces randomness into the MCMC iterations to help avoid local optima and reduces computational costs. At each search position, *S* nearest neighbors (for a small user-configurable constant *S*) with the longest LCS with each estimated haplotype are selected, which are then combined across all positions within the phasing window to obtain the conditioned reference panel. In the TEE setting, the control flow and memory access patterns of this procedure can leak the user’s genotypes, given that the service provider has access to the reference panel. Rigorously preventing this leakage through a direct application of existing mitigations, such as ORAMs, leads to impractical computational overhead, as we show in Figure 4.

TX-Phase introduces an efficient oblivious algorithm for constructing a conditioned reference panel. Our algorithmic advance lies in the development of a compressed PBWT method, which supports the original PBWT routines while operating over the space of unique haplotypes in the compressed panel across contiguous genomic blocks.

The key steps of the algorithm are detailed below. We provide additional detail and pseudocodes for key operations of the compressed PBWT in Supplementary Note 1 and an illustration in Supplementary Figure 5.

#### PBWT on compressed blocks and estimated haplotype insertion

The PBWT algorithm is applied independently to each compressed reference panel block. In contrast to the standard PBWT procedure used in SHAPEIT4, the information generated by PBWT in our approach is encoded in a prefix tree structure, one prefix tree per compressed block. In this structure, each node represents a group of haplotypes that share common prefixes starting from the beginning of the block region, while each leaf corresponds to a unique haplotype within the block. The nodes in each position are lexicographically sorted according to their “reverse prefix” (i.e., suffix in the reverse genomic positions) referred to as the *positional prefix order* in PBWT. *Divergences*, which capture the LCS between adjacent haplotypes in the positional prefix order, are maintained between neighboring nodes. The estimated haplotypes are then partitioned into block regions and inserted into the prefix trees in the correct positional prefix order. While the PBWT on compressed blocks does not involve sensitive data and therefore does not require oblivious algorithms, this haplotype insertion process is performed obliviously to ensure that the paths of the estimated haplotypes within the prefix trees remain confidential. Divergences are updated with each insertion but are kept separate from the prefix trees to distinguish between sensitive and non-sensitive information.

The primary advantage of this prefix tree representation is that it groups haplotypes with common prefixes at every position within the block region. This facilitates the calculation of the block-level LCS between groups of haplotypes and the estimated haplotypes for rapid identification of nearest neighbors.

#### Nearest neighbor candidate searches

To identify candidate haplotypes at each (randomly chosen) search position, the algorithm searches for the nearest nodes to the inserted estimated haplotypes, focusing on those with the longest block-level LCS with the estimated haplotypes. It then combines the haplotypes within these nodes to form the candidate set. This process continues until the candidate set includes at least *S* haplotypes, ensuring that the *S* nearest neighbors—those with the longest global LCS beyond the current block—are among them. The inclusion of the *S* nearest neighbors is guaranteed because a longer block-level LCS indicates a longer global LCS.

The search algorithm leverages the inherent properties of PBWT, which are directly applicable to our prefix tree representation, to perform efficient local searches. In positional prefix order, the nearest nodes are naturally closer together, and the block-level LCS can be computed from the divergences between adjacent nodes. To ensure that the estimated haplotypes’ paths within the prefix tree remain confidential, our oblivious search is performed across all possible paths, with the result for the actual path kept secret.

#### Finding the *S* nearest neighbors

The *S* nearest neighbors among the candidates are ranked based on their global LCS with the estimated haplotypes. Since candidates with shorter block-level LCS also have shorter global LCS, we only need to rank candidates from the node with the shortest block-level LCS to eliminate the excess. To determine the global LCS of these candidates, the global LCS is divided into two non-overlapping regions, which can be combined to assess the global LCS at any position: (1) the global LCS *before* the start of the block region, and (2) the block-level LCS within the block region. Candidates from the node with the shortest block-level LCS share the same (2) by definition, so we only need to compute their (1) for ranking purposes. This computation is done recursively by expanding and connecting their block-level LCS from previous block regions.

To perform this ranking while concealing the paths of the estimated haplotypes within the prefix trees, our approach ranks the top *S* candidates for *every* node based on their (1) and then selects the top candidates from the node with the shortest block-level LCS along the path. This ranking is efficiently carried out using an oblivious merge-sorting algorithm, which aligns naturally with the merge structure of the prefix trees. In the first step, the top *S* candidates for each tree leaf (at the last position in the block region) are identified using an oblivious sorting algorithm. To identify the top *S* candidates for each node in the previous position, a merging step is applied to the child nodes at the current position.

#### Conditioned reference panel construction

After identifying the *S* nearest neighbors at all search positions for both estimated haplotypes, the sets of nearest neighbors are combined using an oblivious bitmap to form the set of conditioned haplotypes. This conditioned set is then used to construct a conditioned reference panel from the compressed reference panel by applying an oblivious filtering algorithm.

### Dynamic Fixed-Point Arithmetic for Accurate Constant-Time Computation

Another key challenge in secure TEE-based phasing is to perform probabilistic calculations accurately without using floating-point numbers, which expose timing side channels [2, 22]. Our phasing algorithm relies heavily on probabilistic calculations involving large matrices and vectors, e.g., in the HMM inference step, where the transition probabilities between adjacent haplotype segments are calculated. Previous works have used fixed-point numbers equipped with constant-time integer operations to address this issue [2, 22]. However, fixed points generally have low precision, which could result in catastrophic underflows, where small probabilities turn into zeros. This in turn reduces the accuracy of phasing, as shown in Figure 5. Although encoding probabilities in log space can offer better precision, this leads to a prohibitive performance overhead in phasing (see Supplementary Table 1).

To address these limitations, TX-Phase introduces *dynamic fixed-point arithmetic*, which can be viewed as a hybrid of fixed-point and floating-point schemes, combining the constant-time properties of the former with the improved precision of the latter. Our key idea, also illustrated in Supplementary Figure 4, is to dynamically maintain an additional scaling factor on a row (or column) basis based on the observation that most of the matrices in the phasing algorithm encode a probability distribution in each row (or column). The scaling helps to retain small numbers in the distribution with minimal impact on the desired calculations, as the scaling factors can be easily accounted for when renormalizing the distribution. We determine the scaling factor as the power of two that would make the largest number in the row (or column) between and 1 after scaling. Multiplying or dividing a vector by this factor can be performed efficiently by applying bit-shift operations to the integers representing the fixed points. Standard arithmetic operations can be performed on our dynamically scaled matrices with additional steps to ensure the scales are consistent. For instance, matrix addition requires that the corresponding rows are shifted to the same scale prior to addition, where the output scale is chosen to minimize precision loss. Element-wise multiplication of two matrices involves multiplying two matrices element-wise, then adding the exponents of the scaling factors of the corresponding rows. Column-wise summation of a matrix requires that all rows be brought to the highest scale within the matrix to minimize precision loss. After each operation on a matrix, we update the scaling factors to maintain the fixed scale of the largest number in each row.

### Other Oblivious Computation Techniques in TX-Phase

TX-Phase incorporates several other transformations applied to the SHAPEIT4 algorithm to ensure that sensitive user data is completely decoupled from observable program behavior with respect to memory access and timing patterns. Sensitive components of the code are systematically detected by our secure typing system, where violations of our security rules are reported as compilation errors. The following summarizes our oblivious computation techniques in specific algorithm components.

#### Initialization

Initialization is a heuristic strategy in the MCMC phasing approach used to estimate the initial phases of the target sample as a starting point for the search. It applies a heuristic function to generate phasing estimates, where each genotype’s phase is determined independently by matching with the nearest neighbors in the PBWT arrays. In TX-Phase, we retrieve the nearest neighbors from the prefix trees, following the same method used for constructing the conditioned reference panel. Additionally, we modify the heuristic function to run in constant time, ensuring that the number of heuristic loops does not depend on sensitive data.

#### Genotype graph construction

The genotype graph structure is determined by the positions of the heterozygous sites in the chromosome. Specifically, the genotype graph nodes in a segment are constructed by iterating all possible haplotypes compatible with three contiguous heterozygous sites in the segment. In TX-Phase, this poses a problem since the positions of the heterozygous sites in the input genotypes are sensitive. The genotype graph structure can reveal where these sites are and the frequency of heterozygous sites. TX-Phase addresses this issue by constructing an oblivious genotype graph that hides the genotype graph segment boundaries. This is accomplished by pretending that every genomic position is a segment boundary and utilizing a secret bitmap of where the true boundaries are when processing the graph.

#### Forward-backward algorithm

The forward-backward algorithm for HMM inference operates by computing the transition probabilities between genotype graph nodes in adjacent segments. However, since the genotype graph in TX-Phase treats all genomic positions as segment boundaries, the forward-backward algorithm is modified to pretend to update the transition probabilities for all supposed segment boundaries while correctly accounting for the true boundaries using a deterministic control flow.

In addition, TX-Phase hides the size of the conditioned reference panel. This indicates how many reference haplotypes are considered the nearest neighbors of the user’s sample, which in principle could reveal private genotypes in certain scenarios. Our oblivious filtering algorithm produces a conditioned reference panel that is padded to a constant size with dummy haplotypes, which are neglected in downstream analysis (i.e., HMM inference) using a bitmap that distinguishes the dummy haplotypes from the real ones.

#### Forward sampling and Viterbi

Both forward sampling and Viterbi rely on a walk in the genotype graph, which can reveal the positions of the heterozygous sites. Following the oblivious solution where all genomic positions are treated as segment boundaries, forward sampling and Viterbi also pretend that all positions are segment boundaries and incorporate them into the algorithms accordingly. However, only the true segment boundaries, kept in the secret map, actually have an effect on the outcome.

#### Pruning

Pruning is an MCMC iteration to increase phasing confidence by merging two adjacent segments whose transition probabilities between top graph nodes are highly likely. To determine which segments are pruned, the original approach in SHAPEIT4 uses a heuristic based on the sum of probabilities of the top nodes and entropy of the segment to rank segment candidates. For oblivious computation, TX-Phase pretends to prune all genomic positions and, instead of ranking the segments, it prunes the segments *greedily* from left to right genomic position if it is determined that the confidence level passes a pruning threshold. Our results show that this modification has a negligible impact on phasing accuracy.

#### Determining phasing genomic windows

In the original approach in SHAPEIT4, the genomic windows for HMM inference are chosen to align with the genotype graph segment boundaries to ensure that the windows always include full segments. However, this presents the same issue where segment boundaries could infer the positions of heterozygous sites. In TX-Phase, we instead determine window sizes by a constant Morgan distance with extra padding genomic positions. The idea is to have the padding wide enough to cover the full segment that would have been truncated by the fixed window size. The size of the padding region can be approximated from the heterozygosity rate, assuming the lower bound is publically known. Our results show that this modification has a negligible impact on phasing accuracy.

### Implementation Details

TX-Phase is implemented in Rust programming language, featuring memory and thread safety and thus preventing the most common security vulnerabilities in software such as memory leaks and race conditions. TX-Phase is deployed in the Gramine framework [23], a Library OS for deploying existing program binaries in an SGX enclave. TX-Phase’s default phasing parameters closely mirror those of SHAPIE4’s to produce the same level of phasing accuracy. The user’s input and output to TX-Phase are in the VCF format. The reference panels are preprocessed into the M3VCF format using Minimac3 [1]. Note that the implementation of TX-Phase is agnostic of the TEE frameworks, so it can likewise be deployed in AMD SEV, Intel TDX, and other SGX frameworks. The strong typing system and constant-time implementation are built on the Rust timing-shield library (https://www.chosenplaintext.ca/open-source/rust-timing-shield/).

### Datasets and Evaluation Settings

To benchmark the accuracy and efficiency of TX-Phase, we utilized the UK Biobank (UKB) dataset (976,754 haplotypes from 488,377 subjects) commonly employed for reference-based phasing benchmarks. We focused our analysis on Chromosome 20, containing 19,959 markers. For cross-validation, we withheld 424 mother-father-child trio samples from the dataset to serve as target samples, with the phasing ground truth determined by trio phasing. Reference panels were constructed by downsampling the full UKB dataset in varying sizes (20K, 40K, 100K, 200K, and 400K haplotypes) to assess the impact of reference panel size on accuracy and efficiency. All types of variants, including single nucleotide polymorphisms and structural variants (insertions and deletions), were included in the benchmarks.

For additional validation, we used the 1000 Genomes Phase 3 (1KG) dataset (5,008 haplotypes from 2,504 subjects) and the Haplotype Reference Consortium (HRC) dataset (54,330 haplotypes from 27,165 subjects) as reference panels, with the Ashkenazi trio from the Genome-in-a-Bottle (GIAB) dataset as the target sample. We focused our analysis on shared markers between reference panels and the target sample on Chromosome 20: 77,593 markers between 1KG and GIAB, and 66,070 between HRC and GIAB. To collect multiple measurements, we run TX-Phase and SHAPEIT4 with 100 random seeds on the same GIAB target sample. Eagle2, being a deterministic algorithm, consistently produces the same output.

The state-of-the-art phasing methods used for comparison include SHAPEIT4 [7] and Eagle2 [6]. SHAPEIT4 serves as a direct comparison as TX-Phase closely follows its phasing algorithm, while Eagle2 represents the tool currently used by the Michigan and TOPMed Imputation Servers. We used the latest versions of SHAPEIT4 (v5.1.1; now part of SHAPEIT5 releases) and Eagle2 (v2.4.1) with default parameters. TX-Phase shares the default SHAPEIT4 parameters, with the addition of a parameter for minimum heterozygous rates in the target samples (used to calculate the size of the phasing window), set to 10% by default.

Reference panels were compressed to M3VCF format for TX-Phase using Minimac3 [1] with default parameters, while remaining in BCF format for SHAPEIT4 and Eagle2. Since TX-Phase does not impute missing genotypes, these were removed from the target samples. To ensure a fair accuracy comparison with SHAPEIT4, which imputes missing genotypes by default, imputed genotypes were removed from its output. For Eagle2, we used the `--noImpMissin` flag to disable imputation.

All experiments were conducted on an Ubuntu server equipped with an Intel Xeon Gold 6426Y processor with SGX version 2, 16 physical CPU cores (Hyperthreading disabled for SGX security), and 128 GB of enclave page cache (EPC). Timing and peak memory usage (based on maximum resident set size) were measured using the `/usr/bin/time` command. Applications running in SGX utilized Gramine [23], a library OS for executing existing binaries within an SGX enclave.

## Data Availability

The UK Biobank genotype array dataset can be accessed via the UKB Research Analysis Platform (RAP) at https://ukbiobank.dnanexus.com. RAP is available to all researchers that are collaborating on an approved in-progress UK Biobank project. The 1000 Genomes Project Phase 3 dataset is publicly available and can be accessed through the International Genome Sample Resource (IGSR) website at http://ftp.1000genomes.ebi.ac.uk/vol1/ftp/phase3/. The Haplotype Reference Consortium dataset is available upon request from the European Genome-Phenome Archive, accession EGAS00001001710. The Genome in a Bottle (GIAB) dataset is publicly available and can be accessed through the National Institute of Standards and Technology (NIST) website at https://ftp-trace.ncbi.nlm.nih.gov/giab/ftp/data/.

## Code Availability

TX-Phase is available under MIT License at https://github.com/hcholab/txphase.

