## Supplementary Materials for "TX-Phase: Secure Phasing of Private Genomes in a Trusted Execution Environment"

##### Supplementary Figures


**Supplementary Figure 1**: TX-Phase's phasing accuracy closely matches SHAPEIT4 and is significantly higher than Eagle2.

**Supplementary Figure 2**: TX-Phase minimizes the computational overhead of TEEs.

**Supplementary Figure 3**: Parallel sample processing with TX-Phase.

**Supplementary Figure 4**: Illustration of our dynamic fixed-point arithmetic.

**Supplementary Figure 5**: Illustration of oblivious phasing via compressed haplotype selection.


##### Supplementary Tables


**Supplementary Table 1**: Our dynamic fixed points enable practical performance of TX-Phase.

**Supplementary Table 2**: Per-operation cost of log-domain fixed-point arithmetic operations.

**Supplementary Table 3**: Notations for our oblivious compressed haplotype selection algorithms.

**Supplementary Table 4**: Operations for our oblivious compressed haplotype selection algorithms.


##### Supplementary Notes


**Supplementary Note 1**: Our oblivious algorithms for compressed haplotype selection.

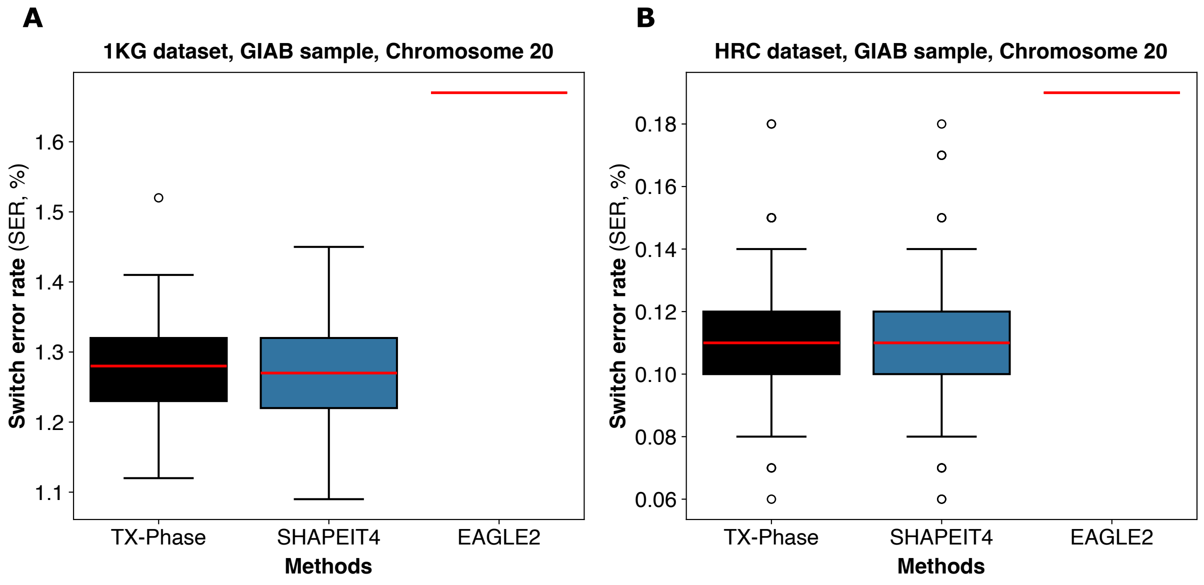


Figure 1: **TX-Phase's phasing accuracy closely matches SHAPEIT4 and is significantly higher than Eagle2.** We evaluated the phasing accuracy of TX-Phase, SHAPEIT4, and Eagle2 on a GIAB Ashkenazi trio (Chromosome 20) using the 1KG (**A**) and HRC (**B**) datasets as reference panels. We repeated the experiment with 100 random seeds for TX-Phase and SHAPEIT4; Eagle2's algorithm is deterministic. Accuracy is measured by switch error rates (SERs). Red horizontal lines indicate the median SER, the boxes represent the interquartile range, and the whiskers extend to the most extreme data points, excluding outliers, which are marked by open circles. TX-Phase SER is not significantly different from SHAPEIT4 when using either the 1KG (two-sided Wilcoxon signed-rank test [WSR] $p = 0.707$) or HRC (WSR $p = 0.443$) reference panels. Eagle2 results in substantially higher SERs in both cases.

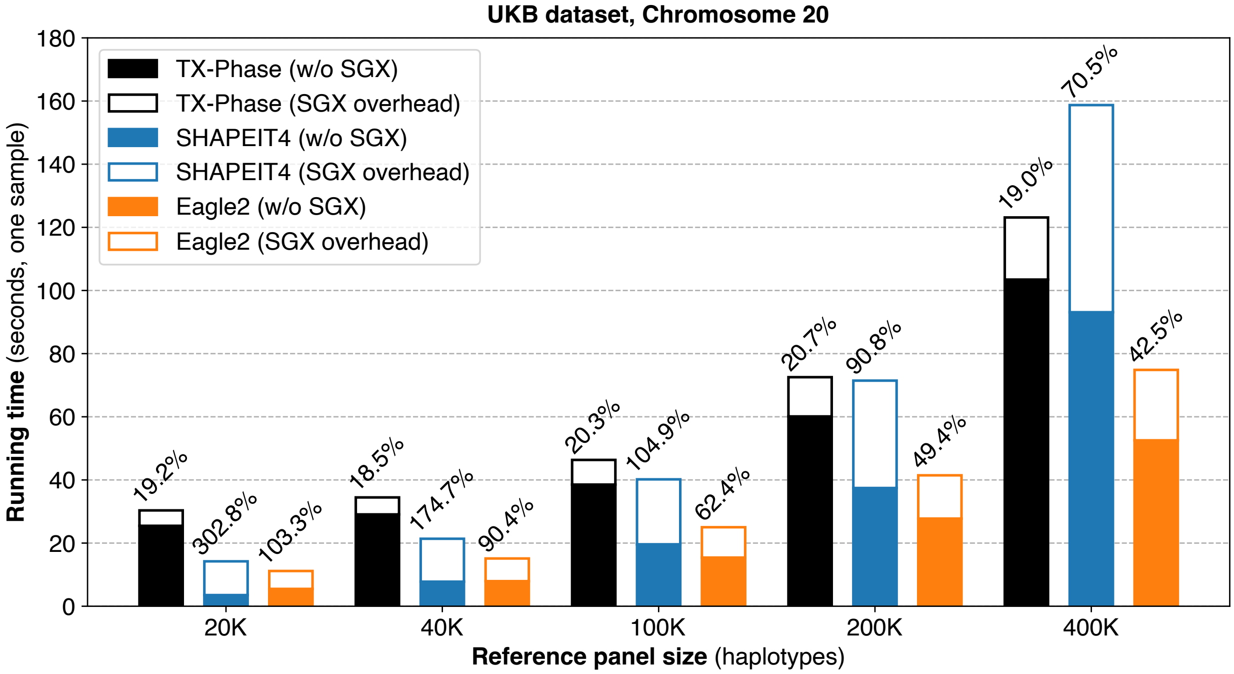


Figure 2: **TX-Phase minimizes the computational overhead of TEEs.** We conducted experiments to compare the overheads of running TX-Phase, SHAPEIT4, and Eagle2 in an SGX enclave compared to outside of SGX. The reported running times are for phasing one target sample (Chromosome 20) with different UKB reference panels. The numbers shown on top of the bars indicate the overhead as a fraction of the baseline running time without SGX. The results demonstrate that TX-Phase's overheads in an SGX enclave are the lowest and consistent at around 20%. In contrast, the overheads are substantially larger in other methods and reduce as the reference panel size grows.

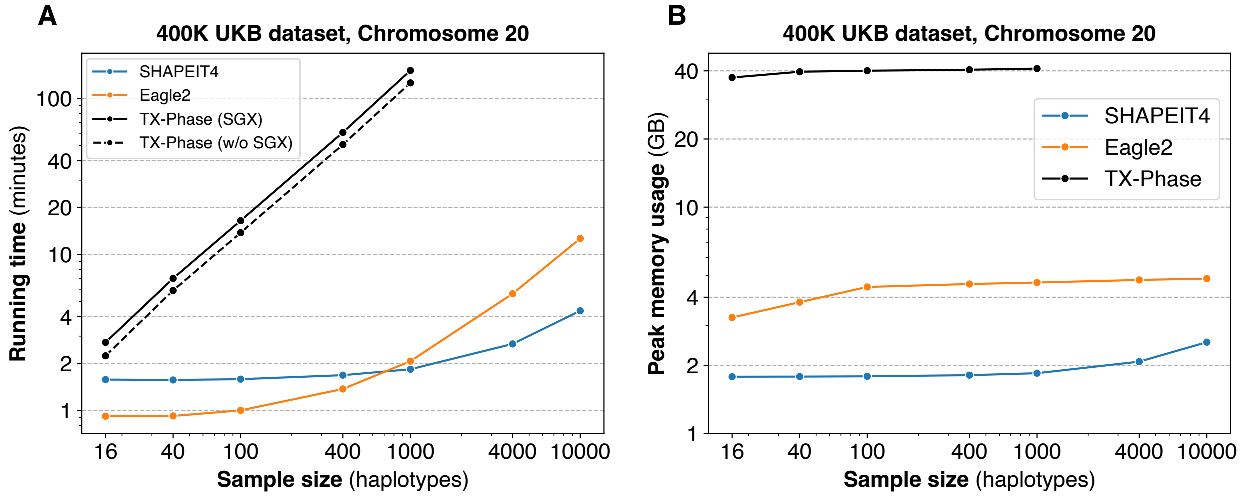


Figure 3: **Parallel sample processing with TX-Phase.** We evaluated the running times of TX-Phase, SHAPEIT4, and Eagle2 for processing multiple samples using a 16-core CPU. TX-Phase is parallelized to process samples in batches of 16 simultaneously. SHAPEIT4 and Eagle2 have native support for multi-threading and incorporate additional algorithmic modifications for handling multiple samples. We measured the running time (**A**) and peak memory usage (**B**) of each method on the 400K haplotype UKB dataset (Chromosome 20) with varying numbers of target samples. Additionally, we measure the running time of TX-Phase without SGX (**A**) to demonstrate the minimal SGX overhead of our method. Although existing tools perform better on large datasets due to additional optimizations for the multi-sample setting, these results confirm that TX-Phase can be easily parallelized to obtain practical performance on moderately sized datasets without incurring additional overheads. For example, we estimate that running TX-Phase on a Microsoft Azure DC48s v3 server (48 CPU cores, 384 GB memory) can phase 5,000 whole-genome samples (805,426 markers in the SNP array) in 5.3 days while utilizing 162 GB of memory, with a total cost of $586 (at $4.608 per hour) based on cost estimates from `https://azure.microsoft.com/en-us/pricing/calculator`. The running time can be further reduced by leveraging more computational resources or incorporating multi-sample optimizations into TX-Phase, which is a meaningful direction for future work. Note that phasing services are mainly intended for target datasets of small to moderate sizes since reference-based phasing offers limited benefit over cohort-based phasing for large cohorts [1].

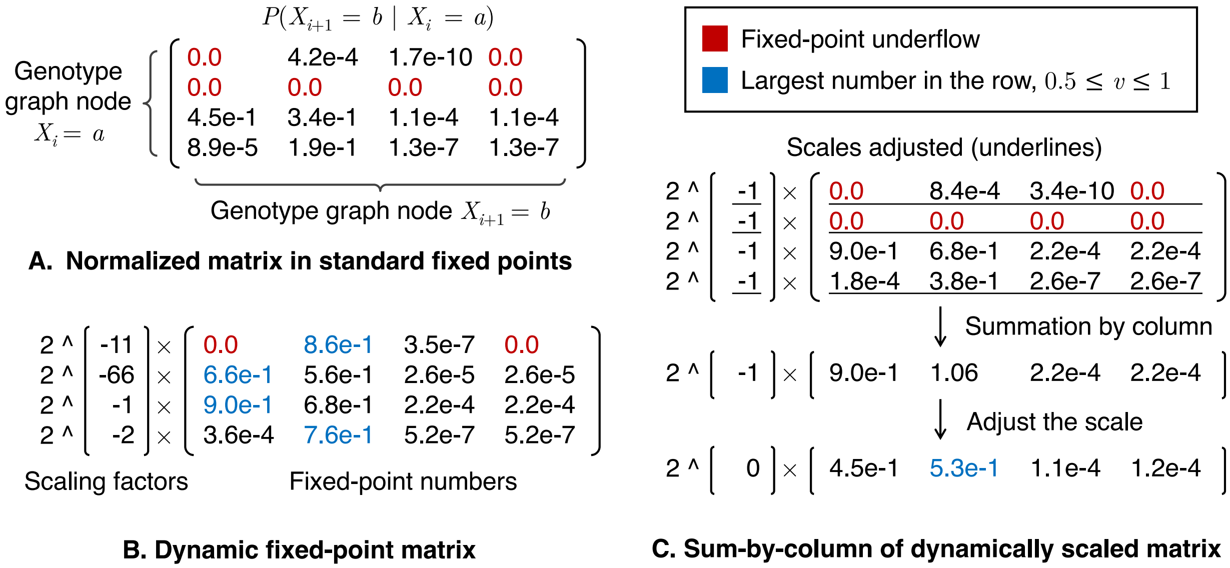


Figure 4: **Illustration of our dynamic fixed-point arithmetic.** (**A**) A transition probability matrix for $P(X_{i+1} = a \| X_i = b)$, where $X_i$ is a genotype graph node at position $i$ and $X_{i+1}$ at $i + 1$, is shown using standard fixed points. Even if the matrix is normalized, probabilities in an entire row can underflow (represented in red), resulting in an accuracy loss. (**B**) The dynamic scaling technique is applied to the matrix, where the scaling factors in powers of two are maintained per row (or column) so that the largest probability in each row (or column) remains between 0.5 and 1.0 (represented in blue). (**C**) A sum-by-column operation is demonstrated on the dynamically scaled matrix. First, the rows with smaller scales (marked with underlines) are adjusted to match the scale of the row with the largest scale to minimize precision loss. This may still result in underflows in some rows but the overall accuracy of the final sum is better preserved. Lastly, the matrix is summed by column, and the scale is readjusted.

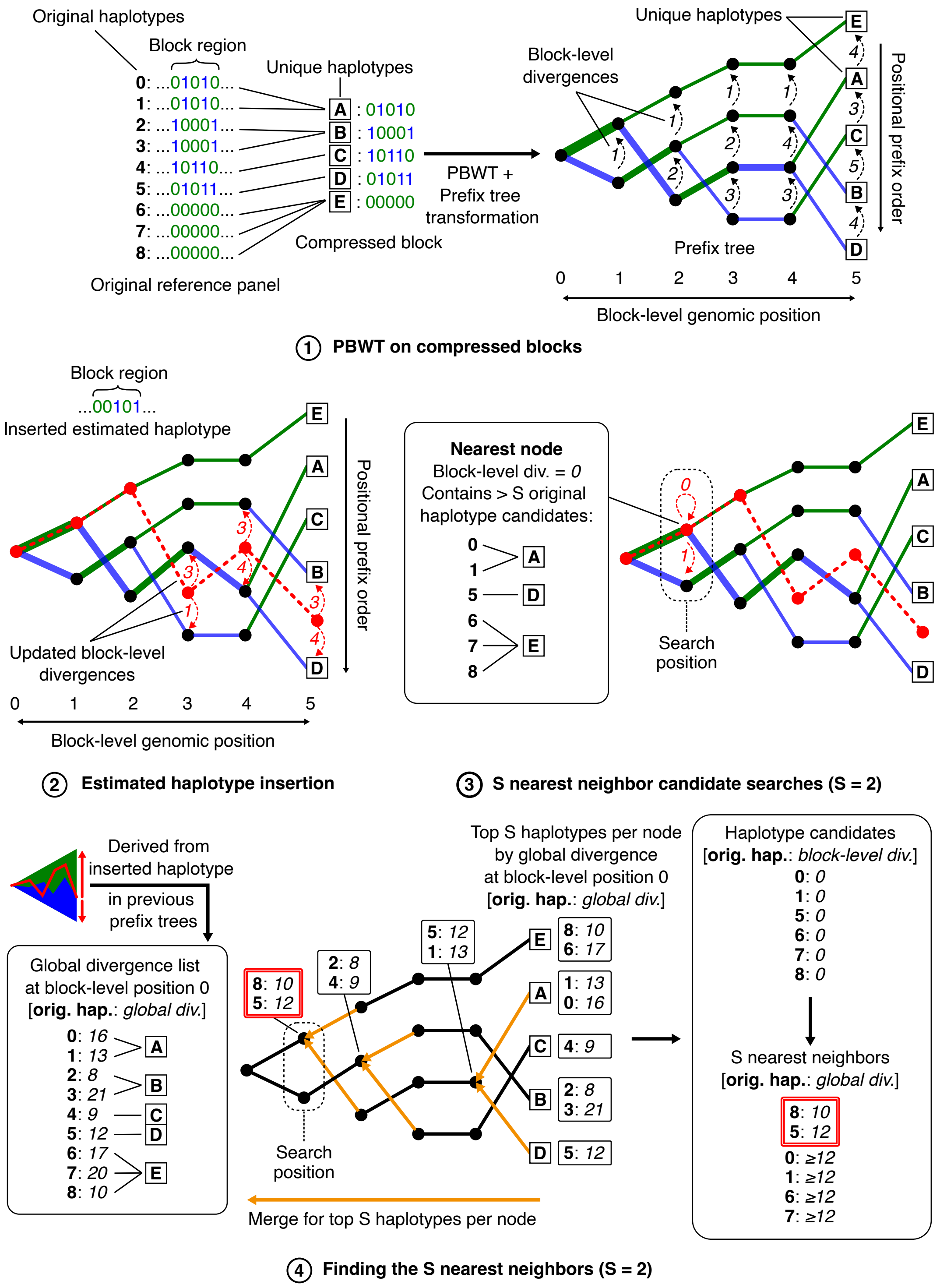


(See caption on next page.)

Figure 5: **Illustration of oblivious phasing via compressed haplotype selection.** We illustrate the workflow of our oblivious compressed haplotype selection, described in detail in Supplementary Note 1. (**1**) Each compressed reference panel block is transformed via PBWT into a prefix tree representation (Algorithm 1). In this tree, each node represents a group of haplotypes that share a common prefix, starting from the root at the beginning of the block region. Each leaf corresponds to a unique haplotype within the block. The nodes are ordered by positional (reverse) prefixes, and block-level divergences (i.e., divergences within the block region) are maintained between adjacent nodes. The 0-edges are shown in green, and the 1-edges in blue, with edge thickness corresponding to the number of haplotypes following that path. (**2**) The portion of the estimated haplotype within the block region is inserted into the tree, while keeping the inserted information separate (Algorithm 2). This includes the updated block-level divergences and positional prefix orders that define the path through the tree. (**3**) At a search position, neighboring nodes with the smallest block-level divergence relative to the estimated haplotype are searched to construct the haplotype candidate set. This set includes original haplotypes along the paths of these nodes, and nodes are searched until at least $S$ original haplotypes with the smallest block-level divergence are identified (Algorithm 3). The candidate set includes the $S$ nearest neighbors because smaller block-level divergences imply smaller global divergences. In this example, where $S = 2$, the candidate set contains six original haplotypes in the nearest node. However, because their block-level divergence is 0, their global divergences are unknown, leaving the order ambiguous. Finding the two nearest neighbors requires further ranking in the next step. (**4**) In our oblivious solution, all original haplotypes within each node are ranked by their global divergences relative to the estimated haplotype at the starting position of the block for the top $S$ haplotypes. The global divergence list can be recursively computed from the estimated haplotype inserted in previous prefix trees (Algorithm 6). By Equation 3, knowing the global divergence at the block's starting position is sufficient to calculate the global divergence at any position within the block, assuming a block-level divergence of 0 (which is the case in this example). The rank of each node is computed using an oblivious merge-sorting algorithm (Algorithm 4): the rank of each leaf node is determined by obliviously sorting the original haplotypes in the node by their divergences from the global divergence list for the top $S$ haplotypes. The rank of each node at higher levels is calculated by merging the ranks of nodes from the level below for the top $S$ haplotypes, following the prefix tree structure. Once the ranks are determined for all nodes, the ranks at the search position are used to resolve the order of haplotype candidates, ultimately determining the $S$ nearest neighbors (Algorithm 5).

| Fixed-point arithmetic | Dynamic scaling | | Log domain | |
|---|---|---|---|---|
| | # function calls | Approx. time | # function calls | Approx. time |
| Addition | $2.9 \times 10^{10}$ | 19 s | $1.1 \times 10^{12}$ | 17.5 m |
| Subtraction | $3.1 \times 10^{6}$ | $< 1$ s | $1.2 \times 10^{11}$ | 2.4 m |
| Multiplication | $3.6 \times 10^{9}$ | 5 s | $9.8 \times 10^{11}$ | 21.1 m |
| Division | $6.7 \times 10^{5}$ | $< 1$ s | - | - |
| | **Total:** | 24 s | **Total:** | 41.0 m |

Table 1: **Our dynamic fixed points enable practical performance of TX-Phase.** We compare the runtime performance between dynamic fixed-point arithmetic techniques, introduced in this work, and the log-domain fixed-point arithmetic techniques from prior work [2]. The comparison is based on a one-sample run on Chromosome 20 using the 400K UKB reference panel. Fixed-point division is implemented in constant time using a bit-wise long division algorithm. The performance of each scheme is measured by the number of function calls to the underlying standard fixed-point arithmetic operations during the run. We then estimate the required runtime by multiplying the number of operations with the observed per-operation costs based on a micro-benchmark, provided in Supplementary Table 2. 's' stands for seconds, and 'm' stands for minutes. The results show that dynamic fixed-point techniques lead to over 100x speedup for TX-Phase compared to the existing techniques.

| | # of function calls to standard fixed-point operations | | |
|---|---|---|---|
| | Addition | Subtraction | Multiplication |
| Log-domain addition | 55 | 6 | 51 |
| Log-domain subtraction | 54 | 5 | 51 |
| Log-domain multiplication | 1 | - | - |
| Log-domain division | - | 1 | - |

Table 2: **Per-operation cost of log-domain fixed-point arithmetic operations.** We analyzed the implementation of log-domain fixed-point arithmetic operations in prior work [2] to assess the costs associated with each arithmetic operation, related to Supplementary Table 1. Specifically, since log-domain fixed points are implemented on top of standard fixed points, we examined the number of function calls made to the underlying standard fixed-point arithmetic operations. This analysis provides insight into how the costs of arithmetic operations multiply when performed in the log domain, in contrast to our dynamic fixed-point arithmetic, where each operation calls exactly one underlying standard fixed-point operation. The results show that log-domain addition and subtraction require a significant number of function calls due to their piecewise polynomial approximation implementations. Consequently, using log-domain fixed points would impose a prohibitively high computational cost for HMM inference in TX-Phase, as shown in Supplementary Table 1. On the other hand, log-domain multiplication and division translate to exactly one addition and one subtraction, respectively, in the normal domain.

| **Notation** | **Definition** |
|---|---|
| $N$ | The number of haplotypes in the original reference panel. |
| $U^b$ | The number of unique haplotypes in the compressed reference panel block $b$. |
| $H^b$ | Compressed reference panel block $b$, where $H^b_{\mathsf{id}}$ denotes unique haplotype $\mathsf{id}$ in the block, and $H^b_{\mathsf{id}}[i]$ denotes the allele of unique haplotype $\mathsf{id} \in \{0, \ldots, U^b - 1\}$ at genomic position $i \in \{0, \ldots, l^b - 1\}$ within the block region. $l^b$ denotes the length of the block region in the number of genomic positions. |
| $l^b$ | The length of the block region $b$ in the number of genomic positions. |
| $B^b$ | The global genomic position (i.e., the position in the entire sequence) at the start of block $b$. Note that $B^b + l^b = B^{b+1}$ |
| $\mathsf{trie}^b_i$ | Prefix tree level $i$ of block region $b$, where $i = 0$ indicates the root level and $i = l^b$ the leaf level. $\mathsf{trie}^b_i$ is a list of tree nodes. |
| $\mathsf{node}$ | A prefix tree node, consisting of the 0-edge ($\mathsf{node}.0$), 1-edge ($\mathsf{node}.1$), the divergence with the node above it ($\mathsf{node.div}$), and the set of unique haplotypes in the node's path ($\mathsf{node.haps}$). |
| $h^b$ | An estimated haplotype of the target sample within block $b$'s region. $h^b[i]$ denotes the allele at block-level genomic position $i \in \{0, \ldots, l^b - 1\}$. |
| $\mathsf{inst}^b_i$ | An haplotype insertion struct at block-level position $i$ in block $b$, marking the insertion of allele $h^b[i-1]$ into the prefix tree level $\mathsf{trie}^b_i$. $\mathsf{inst}^b_i.\mathsf{ord}$ is the order of the inserted haplotype in $\mathsf{trie}^b_i$, where $\mathsf{trie}^b_i[\mathsf{inst}^b_i.\mathsf{ord}]$ indicates the node immediately below the inserted haplotype. $\mathsf{inst}^b_i.\mathsf{div}$ is the block-level divergence between the inserted haplotype and the node immediately above it, $\mathsf{trie}^b_i[\mathsf{inst}^b_i.\mathsf{ord} - 1]$, and $\mathsf{inst}^b_i.\mathsf{div}'$ is the block-level divergence between the inserted haplotype and the node immediately below it, $\mathsf{trie}^b_i[\mathsf{inst}^b_i.\mathsf{ord}]$. |
| $C^b_i$ | The nearest-neighbor candidate list at position $i$ in block $b$, where $j \in C^b_i$ represents node $\mathsf{trie}^b_i[j]$, and $j$ is ordered (ascending) in $C^b_i$ by block-level divergence between node $\mathsf{trie}^b_i[j]$ and estimated haplotype $h^b$ at position $i$. |
| $D_p$ | List of global divergence (i.e. divergence in the entire sequence), where for each original haplotype $\mathsf{id} \in \{0, \ldots, N-1\}$, $D_p[\mathsf{id}]$ is the global divergence between original haplotype $\mathsf{id}$ and an estimated haplotype at position $p$. |
| $\tilde{D}^b_i$ | The block-level divergence (i.e. divergence within a block region) list for block $b$, where for each original haplotype $\mathsf{id} \in \{0, \ldots, N-1\}$, $\tilde{D}^b_i[\mathsf{id}]$ is the block-level divergence between original haplotype $\mathsf{id}$ and an estimated haplotype in the block $h^b$ at block-level position $i$. |
| $R^b_i$ | The rank list of nodes in prefix tree $\mathsf{trie}^b_i$, where $R^b_i[j]$ is the rank of node $\mathsf{trie}^b_i[j]$. Rank is determined by global divergence of original haplotypes. |

Table 3: **Notations for our oblivious compressed haplotype selection algorithms.** Supporting material for Supplementary Note 1.

| **Operations** | **Definition** |
|---|---|
| $L[i:j]$ | Sublist of list $L$ between position $i$ and $j$ (inclusive) |
| $\mathsf{prefix}(\mathsf{node})$ | The prefix of the unique haplotypes in $\mathsf{node}$'s path in the prefix tree. Namely, suppose that $\mathsf{node} \in \mathsf{trie}_i^b$, then $\mathsf{prefix}(\mathsf{node}) := H_{\mathsf{id}}^b[0:i-1]$ for any $\mathsf{id} \in \mathsf{node.haps}$. |
| $L \parallel i$ | Concatenate list $L$ with item or list $i$. |
| **for** $i \in L$ **do** | Iterate all items $i$ in list $L$ in the designated order or set $L$ in an artibary order. |
| $\mathsf{imap}^b(\mathsf{id})$ | Map unique haplotype $\mathsf{id} \in \{0, \ldots, U^b\}$ to the set of original haplotypes (by their IDs) that are compressed to it in block $b$'s region. |
| $\mathsf{Imap}^b(I)$ | Map unique haplotype set $I$ to the combined set of original haplotypes (by their IDs) that are compressed to each unique haplotype in $I$ in block $b$'s region, i.e., $\mathsf{Imap}^b(I) := \bigcup_{\mathsf{id} \in I} \mathsf{imap}^b(\mathsf{id})$. |

Table 4: **Operations for our oblivious compressed haplotype selection algorithms.** Supporting material for Supplementary Note 1.

### Supplementary Note 1: Our Oblivious Algorithms for Compressed Haplotype Selection

This section provides additional details and pseudocodes for compressed haplotype selection in TX-Phase, introduced in Methods (see Accelerating Oblivious Phasing via Compressed Haplotype Selection). The goal of this task is to select a small subset of haplotypes, referred to as *nearest neighbors*, from the reference panel based on their similarity to the phasing estimates. Similarity is determined by identifying the longest shared identity-by-state (IBS) segments. To efficiently identify these haplotypes without exposing the phasing estimates through side channels, our algorithms improve upon the positional Burrows-Wheeler transform (PBWT) [3] used in SHAPEIT4 [4].

In the PBWT-based search, random genomic positions in the sequence are chosen to locate the longest common suffix (LCS) between the reference haplotypes and the phasing estimates, representing the longest IBS segments. For each selected position, the $S$ nearest neighbors with the longest LCS relative to the phasing estimates are identified. To achieve this efficiently, the haplotypes in the reference panel—encoded as 0's and 1's for reference and alternative alleles, respectively—are iteratively transformed via PBWT from the first to the last genomic position to determine their *positional prefix orders* and *divergences* at each position.

The positional prefix order represents the lexicographic ordering of haplotypes based on their reverse prefix, i.e., the sequence of alleles in reverse genomic order starting from the current position and extending back to the first position. A divergence between two haplotypes is defined as the earliest genomic position where they begin to share a common suffix before the current position, which corresponds to the LCS length between the two haplotypes up to that point. It is important to note that the reverse prefix and the suffix of a haplotype at a given position describe the same contiguous region of the haplotype; however, we refer to the reverse prefix when discussing haplotype ordering in the context of PBWT.

In the original PBWT algorithms, a *divergence array* is maintained for each position, storing the divergences between adjacent haplotypes in the positional prefix order. The positional prefix order and divergence arrays possess key properties that facilitate efficient searches for the $S$ nearest neighbors with the longest LCS: (1) haplotypes with longer LCS (i.e., smaller divergence) are positioned closer together in the positional prefix order, and (2) the divergence between any two haplotypes can be calculated by finding the maximum of all divergences between them in the divergence array.

Our algorithms enhance these PBWT approaches by applying PBWT to compressed reference panels, enabling oblivious and efficient nearest-neighbor searches on large datasets. The core idea is to independently apply PBWT to compressed blocks of the reference panel, generating *block-level* divergences (i.e., divergences within a block). These divergences are used to identify nearest-neighbor candidates at each position with the longest block-level LCS. The global divergences (i.e., divergences across the entire sequence) are then computed among these candidates to identify the nearest neighbors with the longest global LCS at a given position.

We first outline the steps of our algorithmic workflow. Summaries of our notation and key operations are provided in Supplementary Tables 3 and 4, respectively. A graphical illustration of our algorithms is provided in Supplementary Figure 5.

**Step 1: PBWT on compressed blocks and prefix tree transformation (Algorithm 1).** This algorithm transforms a compressed reference panel block via PBWT into a prefix tree representation of PBWT arrays, making haplotype insertion and nearest-neighbor searches both oblivious and efficient. The prefix trees compactly preserve all information from the PBWT arrays, grouping haplotypes with a common prefix into a tree node at each position, allowing searches to be performed on groups of haplotypes.

**Step 2: Estimated haplotype insertion (Algorithm 2).** This algorithm inserts the estimated haplotypes of the target sample for the current Markov Chain Monte Carlo (MCMC) iteration into the prefix trees, without revealing their paths within the trees, as such leakage could allow inference of the target sample.

**Step 3: Nearest-neighbor candidate searches (Algorithm 3).** This algorithm quickly identifies a set of nearest-neighbor candidates that is guaranteed to contain the $S$ nearest neighbors of the estimated haplotype at each search position by leveraging the compact prefix tree structure. The $S$ nearest neighbors are the original haplotypes with the smallest global divergence (i.e., longest LCS) with the estimated haplotype at the search position.

**Step 4: Finding the $S$ nearest neighbors (Algorithms 4 and 5).** The algorithms in this step narrow down the haplotype candidates in the candidate node list to identify the $S$ nearest neighbors. This is achieved by ranking the candidates based on their global divergences relative to the estimated haplotype and eliminating those with the largest global divergences. The process is efficient due to the relatively small size of the candidate node list, which is pre-ordered by block-level divergences. Smaller block-level divergences imply smaller global divergences. Therefore, the primary task of these algorithms is to resolve any ambiguities in the ranking based on block-level divergences by computing global divergences as needed.

**Step 5: Computing global divergence lists (Algorithm 6).** This algorithm computes the global divergence lists at the start of the block for all compressed reference panel blocks. These lists are used by Algorithm 4 to rank haplotypes and narrow down the candidate haplotype set to the $S$ nearest neighbors.

**Step 6: Constructing a conditioned reference panel.** In this final step, the nearest neighbors are combined to form the conditioned haplotype set, which is then used to build the conditioned reference panel for HMM inference.

We now provide the details of each step with pseudocode.

#### Step 1: PBWT on compressed blocks and prefix tree transformation (Algorithm 1)

This algorithm (illustrated in Supplementary Figure 5.1) transforms a compressed reference panel block via PBWT into a prefix tree representation of PBWT arrays, making both haplotype insertion and nearest-neighbor searches oblivious and efficient. The prefix trees compactly preserve all information from the PBWT arrays, grouping haplotypes with a common prefix into a tree node at each position, allowing searches to be performed on groups of haplotypes. This step does not involve the sensitive target sample and thus does not require an oblivious implementation.

Algorithm 1 constructs one prefix tree for each compressed reference panel block with the following structure:

- *Prefix tree layers.* The prefix tree for block $b$ consists of tree levels $\mathsf{trie}_i^b$ for each block-level position (i.e., position within the block region) $i \in \{0, \dots, l^b\}$, where $i = 0$ represents the root level and $i = l^b$ represents the leaf level. Each tree level $\mathsf{trie}_i^b$ contains a list of tree nodes.
- *Nodes.* A tree node $\mathsf{node}$ in tree level $\mathsf{trie}_i^b$ consists of the set $\mathsf{node.haps}$ of unique haplotypes in the block that share a common prefix from the root level to level $i$. Specifically, let $H^b$ represent the compressed reference panel block $b$, $H_{\mathsf{id}}^b$ the unique haplotype $\mathsf{id} \in \{0, \dots, U^b - 1\}$ in block $b$, and $H_{\mathsf{id}}^b[i] \in \{0, 1\}$ the allele at block-level position $i$ of unique haplotype $\mathsf{id}$. Then, for all $\mathsf{id} \in \mathsf{node.haps}$ and for all $\mathsf{node} \in \mathsf{trie}_i^b$, the prefix $\mathsf{prefix}(\mathsf{node}) := H_{\mathsf{id}}^b[0 : i - 1]$ is identical between positions 0 and $i - 1$ (inclusive).
- *Prefix tree node order.* The nodes $\mathsf{node} \in \mathsf{trie}_i^b$ are arranged according to the positional prefix order based on the reverse of the prefix $\mathsf{prefix}(\mathsf{node})$.
- *Edges.* A node $\mathsf{node} \in \mathsf{trie}_i^b$ is connected to child nodes in the next tree level, $\mathsf{trie}_{i+1}^b$, via 0-edge $\mathsf{node}.0$ and 1-edge $\mathsf{node}.1$, representing the branching of 0 and 1 alleles, respectively, at the current position of the unique haplotypes in $\mathsf{node.haps}$. Specifically, if $\mathsf{node}.0 = u$ and $\mathsf{node}.1 = v$, then $\mathsf{trie}_{i+1}^b[u].\mathsf{haps} \subseteq \mathsf{node.haps}$ and $\mathsf{trie}_{i+1}^b[v].\mathsf{haps} \subseteq \mathsf{node.haps}$, such that unique haplotype $\mathsf{id} \in \mathsf{trie}_{i+1}^b[u].\mathsf{haps}$ if and only if $\tilde{H}_{\mathsf{id}}^b[i] = 0$, and $\mathsf{id} \in \mathsf{trie}_{i+1}^b[v].\mathsf{haps}$ if and only if $\tilde{H}_{\mathsf{id}}^b[i] = 1$.
- *Divergences.* Each node $\mathsf{node} \in \mathsf{trie}_i^b$ contains the block-level divergence $\mathsf{node.div}$ with the preceding node in the positional prefix order of $\mathsf{trie}_i^b$.
- *Prefix tree root level.* The root tree level $\mathsf{trie}_0^b$ consists of a singleton list containing the root node, which includes all $U^b$ unique haplotypes, i.e., $\mathsf{node.haps} = \{0, \dots, U^b - 1\}$, and $\mathsf{node.div} = 0$.
- *Prefix tree leaf level.* The leaf tree level $\mathsf{trie}_{l^b}^b$ contains leaf nodes with no children, where each node holds a singleton set of one unique haplotype. Thus, every path in the tree represents a unique haplotype in the block.

Intuitively, the above data structure functions similarly to Algorithm 2 in [3]. However, Algorithm 1 enhances this process by independently applying the transformation to each compressed reference panel block, thereby generating block-level positional prefix orders and divergences. It groups unique haplotypes with a common prefix (i.e., divergence 0) into the same prefix tree node. To construct the full prefix tree, Algorithm 1 is applied iteratively at each level, from $i = 0$ to $i = l^b - 1$.

**Algorithm 1 Build prefix tree level $i+1$ for compressed block $b$.** Given prefix tree level $\mathsf{trie}_i^b$ and compressed block $H^b$, build $\mathsf{trie}_{i+1}^b$ and connect the edges of the nodes in $\mathsf{trie}_i^b$ to their child nodes in $\mathsf{trie}_{i+1}^b$. Save the number of 0-edges in level $i$ to $z_i^b$. $b$ is implied and thus omitted for clarity.

---

**Input:** $\mathsf{trie}_i, H$
**Output:** $\mathsf{trie}_i, \mathsf{trie}_{i+1}, z_i$

```
u ← 0, v ← 0, p ← i + 1, q ← i + 1
Create empty lists L_0 and L_1
for node ∈ trie_i do
    p ← max(p, node.div), q ← max(q, node.div)
    Create empty nodes x and y
    for id ∈ node.haps do
        if H_id[i] = 0 then
            if node.0 does not exist then
                node.0 ← u, x.div ← p, p ← 0
            end if
            x.haps ← x.haps ∪ {id}
        else
            if node.1 does not exist then
                node.1 ← v, y.div ← q, q ← 0
            end if
            y.haps ← y.haps ∪ {id}
        end if
    end for
    if x.haps is not empty then
        L_0[u] ← x, u ← u + 1
    end if
    if y.haps is not empty then
        L_1[v] ← y, v ← v + 1
    end if
end for
trie_{i+1} ← L_0 || L_1
for node ∈ trie_i do
    if node.1 exists then
        node.1 ← node.1 + |L_0|          ▷ Offset 1-edges to the positions after 0-edges
    end if
end for
z_i ← |L_0|                              ▷ Save the number of 0-edges
```

#### Step 2: Estimated haplotype insertion (Algorithm 2)

This algorithm (illustrated in Supplementary Figure 5.2) inserts the estimated haplotypes of the target sample for the current Markov Chain Monte Carlo (MCMC) iteration into the prefix trees, without revealing their paths within the trees, as such leakage could allow inference of the target sample. Although each iteration results in two estimated haplotypes per target sample, the insertion and nearest-neighbor searches are independent for each haplotype. Hence, we treat each haplotype independently for the remaining algorithms.

For each block $b$, let $h^b$ represent the portion of the estimated haplotype within block $b$'s region, and $h^b[i] \in \{0,1\}$ for $i \in \{0,\ldots,l^b-1\}$ denote the alleles of the haplotype within that region. Algorithm 2 inserts $h^b$ into the prefix tree for block $b$, while storing the insertion orders and divergences separately in the haplotype insertion structs, $\mathsf{inst}_i^b$, one for each block-level position $i$. Struct $\mathsf{inst}_i^b$ consists of the following elements: $\mathsf{inst}_i^b.\mathsf{ord}$, representing the order of $h^b$ resulting from inserting allele $h^b[i-1]$ into $\mathsf{trie}_i^b$, where node $\mathsf{trie}_i^b[\mathsf{inst}_i.\mathsf{ord}]$ is defined as the node directly below $h^b$ at position $i$; $\mathsf{inst}_i.\mathsf{div}$, representing the block-level divergence between $h^b$ and the node directly *above* it, $\mathsf{trie}_i^b[\mathsf{inst}_i.\mathsf{ord}-1]$; and $\mathsf{inst}_i^b.\mathsf{div}'$, representing the block-level divergence between $h^b$ and the node directly *below* it, $\mathsf{trie}_i^b[\mathsf{inst}_i^b.\mathsf{ord}]$, with $\mathsf{inst}_i^b.\mathsf{div}'$ updating the value of $\mathsf{trie}_i^b[\mathsf{inst}_i.\mathsf{ord}].\mathsf{div}$ after insertion. At the root level, we define $\mathsf{inst}_0^b.\mathsf{ord} = 0$ and $\mathsf{inst}_0^b.\mathsf{div} = \mathsf{inst}_0^b.\mathsf{div}' = 0$.

Algorithm 2 iteratively inserts allele $h^b[i]$ into prefix tree level $\mathsf{trie}_{i+1}^b$, from block-level position $i=0$ to $i=l^b-1$, generating struct $\mathsf{inst}_{i+1}^b$ for each position. At position $i$, the algorithm scans upwards and downwards from the current haplotype order, $\mathsf{inst}_i^b.\mathsf{ord}$, in $\mathsf{trie}_i^b$ until it identifies the first nodes above and below with an $h^b[i]$-edge. Let the identified node above be at order $u$ and the node below at order $v$, with $h^b[i] = E$ for some allele $E$. Then, the next order of $h^b$, $\mathsf{inst}_{i+1}^b.\mathsf{ord}$, is set to $\mathsf{trie}_i^b[u].E+1$, effectively inserting $h^b$ between node $\mathsf{trie}_i^b[u].E$ and node $\mathsf{trie}_i^b[v].E$ in tree level $\mathsf{trie}_{i+1}^b$, following the positional prefix order.

To compute the block-level divergences $\mathsf{inst}_{i+1}^b.\mathsf{div}$ and $\mathsf{inst}_{i+1}^b.\mathsf{div}'$, the algorithm uses the PBWT property, taking the maximum of divergences between $h^b$ and nodes $u$ and $v$ at position $i$:

$$\mathsf{inst}_{i+1}^b.\mathsf{div} = \max\left(\mathsf{inst}_i^b.\mathsf{div}, \mathsf{trie}_i^b[j-1].\mathsf{div}, \mathsf{trie}_i^b[j-2].\mathsf{div}, \ldots, \mathsf{trie}_i^b[u-1].\mathsf{div}\right) \tag{1}$$

$$\mathsf{inst}_{i+1}^b.\mathsf{div}' = \max\left(\mathsf{inst}_i^b.\mathsf{div}', \mathsf{trie}_i^b[j+1].\mathsf{div}, \mathsf{trie}_i^b[j+2].\mathsf{div}, \ldots, \mathsf{trie}_i^b[v].\mathsf{div}\right), \tag{2}$$

where $j := \mathsf{inst}_i.\mathsf{ord}$.

In our oblivious implementation of Algorithm 2, the **for** loops always run between the lower bound (at 0 or 1) and the upper bound (at $|\mathsf{trie}_i^b|-1$), pretending to compute within the loop to mask the actual value of $\mathsf{inst}_i.\mathsf{ord}$. The **break** statement does not terminate the loop but triggers a condition where computation continues without further updates to the relevant variables.

**Algorithm 2 Insert an estimated haplotype at position $i+1$ within block $b$.** Given haplotype insertion struct $\mathsf{inst}_i^b$, allele $h^b[i]$, tree level $\mathsf{trie}_i^b$, and the number of 0-edges in $\mathsf{trie}_i^b$, $z_i^b$, build struct $\mathsf{inst}_{i+1}^b$. $b$ is implied and thus omitted for clarity.

```
Input:   inst_i, h[i], trie_i, z_i
Output:  inst_{i+1}

Create an empty struct inst_{i+1}
                                        ▷ Compute inst_{i+1}.ord and inst_{i+1}.div below
d ← inst_i.div; found ← false
for j = inst_i.ord − 1 → 0 do                    ▷ Scan upward for the first h[i]-edge
    node ← trie_i[j]
    if h[i] = 0 and node.0 exists then
        inst_{i+1}.ord ← node.0 + 1; inst_{i+1}.div ← d; found ← true
        break
    else if h[i] = 1 and node.1 exists then
        inst_{i+1}.ord ← node.1 + 1; inst_{i+1}.div ← d; found ← true
        break
    end if
    d ← max(d, node.div)                                   ▷ Compute Equation 1
end for
if not found then
    if h[i] = 0 then
        inst_{i+1}.ord ← 0
    else
        inst_{i+1}.ord ← z_i
    end if
    inst_{i+1}.div ← i + 1
end if
                                                  ▷ Compute inst_{i+1}.div′ below
d ← inst_i.div′; found ← false
for j = inst_i.ord → |trie_i| − 1 do             ▷ Scan downward for the first h[i]-edge
    node ← trie_i[j]
    if j > inst.ord then
        d ← max(d, node.div)                               ▷ Compute Equation 2
    end if
    if (h[i] = 0 and node.0 exists) or (h[i] = 1 and node.1 exists) then
        inst_{i+1}.div′ ← d; found ← true
        break
    end if
end for
if not found then
    inst_{i+1}.div′ ← i + 1
end if
```

#### Step 3: Nearest-neighbor candidate searches (Algorithm 3)

This algorithm (illustrated in Supplementary Figure 5.3) quickly identifies a set of nearest-neighbor candidates that is guaranteed to contain the $S$ nearest neighbors of the estimated haplotype at each search position by leveraging the compact prefix tree structure. The $S$ nearest neighbors are the original haplotypes with the smallest global divergence (i.e., longest LCS) with the estimated haplotype at the search position.

In the PBWT-based search, several genomic positions in the sequence are randomly chosen for a nearest-neighbor search. Suppose position $i$ in block $b$ is selected; this algorithm identifies the set of original haplotype candidates that includes all $S$ original haplotypes with the smallest global divergences relative to the estimated haplotype at the given position. Since smaller block-level divergences imply smaller global divergences, these candidate original haplotypes can be found by searching for nodes containing original haplotypes with the smallest block-level divergences with the estimated haplotype until at least $S$ original haplotypes are identified. If the candidate set contains more than $S$ haplotypes, it will be narrowed down to exactly $S$ haplotypes in the next step by their global divergences.

Algorithm 3 represents the candidate set at position $i$ in block $b$ using the candidate node list $C_i^b$, where for all $j \in C_i^b$, $j$ represents node $\mathsf{trie}_i^b[j]$ and is ordered by the smallest block-level divergence between the node (specifically, the node's prefix, $\mathsf{prefix}(\mathsf{trie}_i^b[j])$) and the estimated haplotype $h^b$. The candidate node list forms the candidate set by combining all the original haplotypes along the paths of these nodes. Let $\mathsf{imap}^b(\mathsf{id})$ be the index map between the unique haplotype $\mathsf{id}$ in block $b$ and the set of original haplotypes compressed to it within the block $b$'s region, and define $\mathsf{Imap}^b(\mathsf{node.haps}) := \bigcup_{\mathsf{id} \in \mathsf{node.haps}} \mathsf{imap}^b(\mathsf{id})$. The candidate set is the combined set $\bigcup_{j \in C_i^b} \mathsf{Imap}^b(\mathsf{trie}_i^b[j].\mathsf{haps})$, which includes at least $S$ members.

Intuitively, Algorithm 3 constructs the candidate list $C_i^b$ by performing a local scan for nodes with the smallest block-level divergences relative to the estimated haplotype $h^b$ in the prefix tree. This process leverages the properties of PBWT, where: (1) nodes with the smallest block-level divergences relative to the estimated haplotype $h^b$ are nearest to its order, $\mathsf{inst}_i^b.\mathsf{ord}$, in $\mathsf{trie}_i^b$, and (2) the nodes' block-level divergences with $h^b$ can be dynamically computed from the nodes' divergences, similar to how Equations 1 and 2 calculate the divergence between $h^b$ and its neighboring nodes. This approach allows Algorithm 3 to efficiently scan locally around order $\mathsf{inst}_i^b.\mathsf{ord}$ in $\mathsf{trie}_i^b$ to identify the candidate nodes with the smallest block-level divergences with respect to $h^b$.

In our oblivious implementation of Algorithm 3, we conceal the value of the haplotype insertion struct $\mathsf{inst}_i^b$ by computing candidate lists for all possible values of $\mathsf{inst}_i.\mathsf{ord}$, and then employing an ORAM algorithm (see Methods) to select the candidate list that corresponds to the true value of $\mathsf{inst}_i^b.\mathsf{ord}$.

**Algorithm 3 Search for nearest-neighbor candidates at positions $i$.** Given haplotype insertion struct $\mathsf{inst}_i^b$ for the estimated haplotype $h^b$ and tree level $\mathsf{trie}_i^b$, compute the list $C_i^b$ of candidate nodes with the smallest block-level divergences with $h^b$ in block $b$. $b$ is implied and thus omitted for clarity.

**Input:** $\mathsf{inst}_i, \mathsf{trie}_i$
**Output:** $C_i$

```
Create an empty list C_i
n ← 0                                      ▷ Keep the count of original haplotype candidates
u ← inst_i.ord − 1, v ← inst_i.ord
p ← inst_i.div, q ← inst_i.div'
while n < S do
    if p ≤ q then
        C_i ← C_i || u
        n ← n + |lmap(trie_i[u].haps)|     ▷ Add the node's original haplotypes
        p ← max(p, trie_i[u].div)
        u ← u − 1
    else
        C_i ← C_i || v
        n ← n + |lmap(trie_i[v].haps)|     ▷ Add the node's original haplotypes
        q ← max(q, trie_i[v + 1].div)
        v ← v + 1
    end if
end while
```

#### Step 4: Finding the $S$ nearest neighbors (Algorithms 4 and 5)

The algorithms (illustrated in Supplementary Figure 5.4) in this step narrow down the haplotype candidates in the candidate node list to identify the $S$ nearest neighbors. This is achieved by ranking the candidates based on their global divergences relative to the estimated haplotype and eliminating those with the largest global divergences. The process is efficient due to the relatively small size of the candidate node list, which is pre-ordered by block-level divergences. Smaller block-level divergences imply smaller global divergences. Therefore, the primary task of these algorithms is to resolve any ambiguities in the ranking based on block-level divergences by computing global divergences as needed.

To understand this process, consider the relationship between global divergences and block-level divergences. Let $D_p$ be the *global divergence list* at global position $p$, where $D_p[\mathsf{id}]$ represents the global divergence between the original haplotype $\mathsf{id} \in \{0, \ldots, N-1\}$ and the estimated haplotype at position $p$. Similarly, let $\tilde{D}^b_i$ be the *block-level divergence list* at block-level position $i$ within block $b$, where $\tilde{D}^b_i[\mathsf{id}]$ denotes the block-level divergence between the original haplotype $\mathsf{id} \in \{0, \ldots, N-1\}$ and the estimated haplotype $h^b$ at position $i$. The global divergence $D_p[\mathsf{id}]$ can be decomposed into two components:

1. The global divergence $D_{B^b}[\mathsf{id}]$ at the starting position of block $b$, denoted $B^b$.

2. The block-level divergence $\tilde{D}^b_i[\mathsf{id}]$ at position $i$ within block $b$, where $B^b + i = p$.

Given these components, the global divergence $D_p[\mathsf{id}]$ can be computed using the following equation:

$$D_p[\mathsf{id}] = \begin{cases} D_{B^b}[\mathsf{id}], & \text{if } \tilde{D}^b_i[\mathsf{id}] = 0 \\ B^b + \tilde{D}^b_i[\mathsf{id}], & \text{if } \tilde{D}^b_i[\mathsf{id}] > 0 \end{cases} \quad (3)$$

Intuitively, if $\tilde{D}^b_i[\mathsf{id}] = 0$, it indicates that the LCS between the original haplotype $\mathsf{id}$ and the estimated haplotype spans from block-level position 0 to $i-1$ and may also extend before block $b$'s region. Thus, the global divergence at global position $p$ is the same as the global divergence preceding block $b$'s region, $D_{B^b}[\mathsf{id}]$. On the other hand, if $\tilde{D}^b_i[\mathsf{id}] > 0$, the LCS is contained within block $b$'s region, so the global divergence at position $p$ is updated by offsetting the block-level divergence $\tilde{D}^b_i[\mathsf{id}]$ with the starting position of the block region, $B^b$.

Given Equation 3, we first identify scenarios where the ranking of haplotype candidates in the candidate node list $C^b_i$ is *unambiguous*, and therefore does *not* require computing global divergences to determine the $S$ nearest neighbors:

1. *The candidate set contains exactly $S$ candidates.* Since these candidates are already known to have the smallest block-level divergences, and hence the smallest global divergences, no further action is required.

2. *The last node in the candidate node list contains haplotype candidates with a non-zero block-level divergence.* Specifically, let $j$ be the last node in $C^b_i$. In this case, for all $\mathsf{id} \in \mathsf{Imap}^b(\mathsf{trie}^b_i[j].\mathsf{haps})$, it holds that $\tilde{D}^b_i[\mathsf{id}] > 0$. Since $C^b_i$ is ordered by the smallest block-level divergence between the candidate nodes and the estimated haplotype $h^b$, the haplotype candidates with the largest block-level divergences—and thus the largest global divergences—are located in the last node $j$. As all haplotypes $\mathsf{id} \in \mathsf{Imap}^b(\mathsf{trie}^b_i[j].\mathsf{haps})$ have the same block-level divergence $\tilde{D}^b_i[\mathsf{id}] > 0$ due to their common prefix by definition, we can simply eliminate any haplotypes from this node until only $S$ nearest neighbors remain.

This leaves us with the only scenario where the ranking is ambiguous: *when there is only one node on the candidate node list, and that node contains more than $S$ haplotype candidates with zero block-level divergence.* Specifically, let $j$ be the only node in $C_i^b$. In this case, for all $\mathsf{id} \in \mathsf{Imap}^b(\mathsf{trie}_i^b[j].\mathsf{haps})$, it holds that $\tilde{D}_i^b[\mathsf{id}] = 0$, and $|\mathsf{Imap}^b(\mathsf{trie}_i^b[j].\mathsf{haps})| > S$. Here, the ranking is ambiguous because, according to Equation 3, the global divergence is given by $D_p[\mathsf{id}] = D_{B^b}[\mathsf{id}]$ but $D_{B^b}[\mathsf{id}]$ is unknown. Suppose that $D_{B^b}[\mathsf{id}]$ can be computed (we detail how this is done in the next step), then the haplotype candidates can be ranked by the smallest $D_{B^b}[\mathsf{id}]$ to keep the $S$ nearest neighbors with the smallest global divergence.

The approach of the algorithm in this step is to distinguish between these scenarios and resolve the ambiguity as needed. However, doing this in the oblivious implementation could potentially expose the value of $\tilde{D}_i^b[\mathsf{id}]$ via side channels, and thereby reveal the similarity between the estimated haplotype and the reference haplotypes. Moreover, the choice of which nodes are ranked by global divergences could disclose the path of the estimated haplotype in the prefix trees.

To mitigate these potential issues, our oblivious implementation addresses all scenarios simultaneously by following these two steps: first, in Algorithm 4, haplotypes in *all* nodes are ranked by their global divergence at the starting position of the block, $D_{B^b}[\mathsf{id}]$; then, in Algorithm 5, the resulting ranks along the path of the estimated haplotype are chosen to resolve ranking ambiguities and determine the $S$ nearest neighbors.

In Algorithm 4, our oblivious algorithm efficiently computes the ranks of haplotypes in all nodes using the values of $D_{B^b}[\mathsf{id}]$ via an oblivious merge-sorting algorithm. Intuitively, the algorithm obliviously sorts haplotypes in each leaf node independently. Then, at the next level—from the level above the leaves to the root—the top haplotypes are merged according to the prefix tree structure.

Let $R_i^b$ be the *rank list* at block-level position $i$ in block $b$. The *rank* $R_i^b[j]$ of node $\mathsf{trie}_i^b[j]$ is defined as the list of original haplotypes $\mathsf{id} \in \mathsf{Imap}^b(\mathsf{trie}_i^b[j].\mathsf{haps})$, sorted by $D_{B^b}[\mathsf{id}]$ in ascending order, with $|R_i^b[j]| = \min(S, |\mathsf{Imap}^b(\mathsf{trie}_i^b[j].\mathsf{haps})|)$. The oblivious merge-sorting algorithm proceeds as described in Algorithm 4.

Then, in Algorithm 5, the ranks are combined to select the $S$ nearest neighbors based on the candidate node list. This algorithm combines ranks, irrespective of whether the haplotype order is ambiguous, without affecting correctness, since the internal order within the final set of $S$ nearest neighbors is insignificant.

The oblivious implementation of Algorithm 5 is integrated with the oblivious implementation of Algorithm 3, where the candidate lists $C_i^b$ for all possible values of $\mathsf{inst}_i^b.\mathsf{ord}$ are computed. Then, Algorithm 5 is applied to all the resulting $C_i^b$, and ORAM is used to select the $S$ nearest neighbors for the correct $C_i^b$ at the end.

**Algorithm 4 Compute ranks.** Given the prefix tree for block $b$, $\mathsf{trie}_i^b$ for all $i \in \{1, \ldots, l^b\}$, and the global divergence list $D_{B^b}$ at the starting position of block $b$, $B^b$, compute the rank lists $R_i^b$ for all $i \in \{1, \ldots, l^b\}$.

**Input:** $\mathsf{trie}_i^b$ for all $i \in \{1, \ldots, l^b\}, D_{B^b}$
**Output:** $R_i^b$ for all $i \in \{1, \ldots, l^b\}$

```
Create empty lists R_i^b for all i ∈ {1, ..., l^b}
for i = l^b → 1 do
    for j = 0 → |trie_i^b| − 1 do
        G ← lmap^b(trie_i^b[j].haps)
        s ← min(S, |G|)
        if i = l^b then                                          ▷ Initialization step
            R_i^b[j] ← top s id ∈ G, sorted by the smallest D_{B^b}[id]
        else                                                     ▷ Merging step
            u ← trie_i^b[j].0, v ← trie_i^b[j].1
            r_0 ← R_{i+1}^b[u], r_1 ← R_{i+1}^b[v]
            R_i^b[j] ← top s id across r_0 and r_1, sorted by the smallest D_{B^b}[id]
        end if
    end for
end for
```

**Algorithm 5 Find the $S$ nearest neighbors.** Given the candidate list $C_i^b$ and the rank list $R_i^b$, find the $S$ nearest neighbor set $I_i^b$ of the estimated haplotype. $b$ is implied and thus omitted for clarity.

**Input:** $C_i, R_i$
**Output:** $I_i$

```
Create an empty set I_i
for j ∈ C_i do
    s ← min(S − |I_i|, |R_i[j]|)
    G ← (R_i[j])[0 : s − 1] as a set
    I_i ← I_i ∪ G
end for
```

#### Step 5: Computing global divergence lists (Algorithm 6)

This algorithm computes the global divergence lists $D_{B^b}$ at the start of block $b$, $B^b$, for all compressed reference panel blocks. These lists are used by Algorithm 4 to rank haplotypes and narrow down the candidate haplotype set to the $S$ nearest neighbors.

Algorithm 6 leverages the observation from Equation 3 that the global divergence list $D_{B^{b+1}}$ at the start of block $b+1$ can be recursively derived from the global divergence list $D_{B^b}$ at the start of block $b$ and the block-level divergence list $\tilde{D}^b_{l^b}$ at the last position of block $b$ (where $B^b + l^b = B^{b+1}$), starting from the initial list $D_0 = \{0\}^N$. The value of $\tilde{D}^b_{l^b}[\mathsf{id}]$ is calculated by the block-level divergence between the estimated haplotype $h^b$ and node $\mathsf{node} \in \mathsf{trie}^b_{l^b}$, where $\mathsf{id} \in \mathsf{Imap}^b(\mathsf{node.haps})$. This is done by utilizing the PBWT to compute divergences dynamically: by scanning up and down the prefix tree level $\mathsf{trie}^b_{l^b}$ and determining the maximum divergence between the estimated haplotype at $\mathsf{inst}^b_{l^b}.\mathsf{ord}$ and the target node $\mathsf{node}$ such that $\mathsf{id} \in \mathsf{Imap}^b(\mathsf{node.haps})$.

**Algorithm 6 Compute global divergent lists.** Given the global divergence list $D_{B^b}$, at the starting position of block $b$, prefix tree leaf level $\mathsf{trie}_{l^b}^b$ at the last position of block $b$, and the haplotype insertion struct $\mathsf{inst}_{l^b}^b$ at the last position of block $b$, compute $D_{B^{b+1}}$ at the starting position of block $b+1$.

```
Input:    D_{B^b}, trie^b_{l^b}, inst^b_{l^b}
Output:   D_{B^{b+1}}
                                                  ▷ Compute the block-level divergence list
D~^b_{l^b} ← {0}^N
d ← inst^b_{l^b}.div
for j = inst^b_{l^b}.ord − 1 → 0 do                                         ▷ Scan upward
    for id ∈ Imap^b(trie^b_{l^b}[j].haps) do      ▷ Note that |trie^b_{l^b}[j].haps| = 1 by definition
        D~^b_{l^b}[id] ← d
    end for
    d ← max(d, trie_{l^b}[j].div)
end for
d ← inst_{l^b}.div'
for j = inst_{l^b}.ord → |trie_{l^b}| − 1 do                              ▷ Scan downward
    if j > inst.ord then
        d ← max(d, trie_i[j].div)
    end if
    for id ∈ Imap^b(trie^b_{l^b}[j].haps) do
        D~^b_{l^b}[id] ← d
    end for
end for
                                                ▷ Compute the global divergence list
D_{B^{b+1}} ← {0}^N
for j = 0 → N − 1 do
    if D~^b_{l^b}[j] = 0 then                                          ▷ Compute Equation 3
        D_{B^{b+1}}[j] ← D_{B^b}[j]
    else
        D_{B^{b+1}}[j] ← B^b + D~^b_{l^b}[j]
    end if
end for
```

#### Step 6: Constructing a conditioned reference panel

This is the final step of the oblivious compressed haplotype selection algorithms, where the nearest neighbors are combined to form the conditioned haplotype set, which is then used to build the conditioned reference panel for HMM inference. To construct the conditioned haplotype set, each of the two estimated haplotypes of the target sample undergoes independent $S$ nearest neighbor searches at the same randomly chosen positions. All the $S$ nearest neighbors of both estimated haplotypes at these positions are combined to form a unified conditioned haplotype set. The haplotype IDs in this set are used to look up and copy the corresponding unique haplotypes from the compressed reference panel into the conditioned reference panel. The efficiency gain of this approach is a result of the smaller memory footprint of the compressed reference panel, allowing for faster lookups and copying.

Our oblivious implementation of this step ensures that both the selected haplotypes and the size of the conditioned haplotype set remain hidden. This involves two key operations: first, obliviously computing the set union of all the nearest neighbors, and second, obliviously filtering the compressed reference panel based on the conditioned haplotype set.

To compute the set union obliviously, we create an oblivious bitmap, where each bit indicates whether the original haplotype $\mathsf{id} \in \{0, \ldots, N-1\}$ is part of the conditioned haplotype set. To incorporate a new $S$-nearest-neighbor set into the conditioned haplotype set, we iterate through all haplotypes in the $S$-nearest-neighbor set and use ORAM access to look up the haplotypes and flip the corresponding bits in the oblivious bitmap. Once the bitmap is fully updated, each original haplotype ID is mapped to its corresponding unique haplotype ID for each compressed block. Using ORAM access, the unique haplotype IDs are then used to look up and copy the unique haplotypes from the compressed reference panel blocks to build the conditioned reference panel. Given the small size of the array due to bit-packing, we use a linear scan ORAM, which leads to better concrete efficiency than other more complex ORAM methods.
